# Glycan heterogeneity as a cause of the persistent fraction in HIV-1 neutralization

**DOI:** 10.1101/2023.08.08.552396

**Authors:** Rajesh P. Ringe, Philippe Colin, Gabriel Ozorowski, Joel D. Allen, Anila Yasmeen, Gemma E. Seabright, Jeong Hyun Lee, Aleksandar Antanasijevic, Kimmo Rantalainen, Thomas Ketas, John P. Moore, Andrew B. Ward, Max Crispin, P. J. Klasse

## Abstract

Neutralizing antibodies (NAbs) to multiple epitopes on the HIV-1 envelope glycoprotein (Env) have been isolated from infected persons. The potency of NAbs is more often measured than the size of the persistent fraction *o*f infectivity at maximum neutralization, which may also influence preventive efficacy by active or passive immunization and the therapeutic outcome of the latter. HIV-1 CZA97.012, a clone of a Clade C isolate, is neutralized to ∼100% by many NAbs. But here NAb PGT151, directed to a fusion-peptide epitope, was shown to leave a persistent fraction of 15%. NAb PGT145, ligating the Env-trimer apex, was less potent but more effective. We sought explanations of the different persistent fractions by depleting pseudoviral populations of the most PGT151- and PGT145-reactive virions. Thereby, neutralization by the non-depleting NAb increased; it decreased by the depleting NAb. Furthermore, depletion by PGT151 increased sensitivity to autologous neutralization by sera from rabbits immunized with soluble native-like CZA97.012 trimer: substantial persistent fractions were reduced. NAbs in these sera target epitopes comprising residue D411 at the V4-β19 transition in a defect of the glycan shield on CZA97.012 Env. Affinity-fractionated soluble native-like CZA97.012 trimer showed commensurate antigenic differences in analyses by ELISA and surface plasmon resonance. We then demonstrated glycan differences between PGT151- and PGT145-purified trimer fractions by mass spectrometry, providing one explanation for the differential antigenicity. These differences were interpreted in relation to a new structure at 3.4-Å resolution of the soluble CZA97.012 trimer determined by cryo-electron microscopy. PGT151-purified trimer showed a closed conformation, refuting apex opening as the cause of reduced PGT145 binding. The evidence suggests that differences in binding and neutralization after trimer purification or PV depletion with PGT145 or PGT151 are caused by variation in glycosylation, and that some glycan variants confer antigenic heterogeneity through direct effects on antibody contacts, whereas others act allosterically.

**Author Summary:** Neutralizing antibodies block the entry of HIV-1 into cells and protect against HIV-1 infection in animal models. Therefore, a goal of vaccination is to elicit antibodies that potently neutralize most HIV-1 variants. Such antibodies suppress virus levels when given to HIV-1-infected patients. Their potency is often measured as the concentration that gives 50% or 80% neutralization. But higher degrees of neutralization are needed to protect an organism from infection. And for some antibodies a ceiling is reached, so that even with increased concentrations a constant fraction of infectious virus persists. We studied the carbohydrate moieties on the envelope glycoprotein, which is the sole target for neutralizing antibodies, of one HIV-1 isolate of the most widespread subtype, Clade C, prevalent in Africa and Asia. We show how differences in carbohydrates can contribute to persistent infectivity, because distinct carbohydrates fit different antibodies. With a new three-dimensional structure of the entry-mediating protein from the Clade-C isolate, we illustrate that some carbohydrate differences occur exactly where the antibodies bind, whereas others are located elsewhere and can act indirectly. When we combined two neutralizing antibodies the persistent infectivity shrank. Our results reinforce the need for multiple specificities of neutralizing antibodies in prevention and therapy.

## INTRODUCTION

Neutralizing antibodies (NAbs) are the best correlate of protection against several viral infections (1–7). Broadly active NAbs (bNAbs) protect against HIV-1 acquisition in animal models, but eliciting them through active immunization with their sole target, the viral envelope glycoprotein (Env), has proved elusive (8–17). In contrast, therapeutic passive immunization of HIV-1-infected people by the administration of bNAbs already shows promise in clinical trials (12, 18–20). And preventive passive immunization specifically blocks acquisition of isolates that are sensitive to the administered bNAb (21, 22).

When bNAbs are evaluated, much focus is on their breadth of action against panels of relatively neutralization-resistant isolates and on their efficiency or potency, *i.e.*, the concentration of pure bNAb that gives, *e.g*., 50% or 80% neutralization. Arguably, however, their efficacy, *i.e.*, their maximum extent of neutralization of an individual isolate can also influence their preventive and therapeutic effect.

The converse of the maximum of virus neutralization, known as the persistent fraction of infectivity (PF), has a long and contentious history in virology (1, 23, 24). The PF can be detected as the neutralization by NAb at increasing concentration reaches a maximum below 100% (1, 23–25). Generally, the persistent virus is not genetically resistant, but when propagated *de novo* and re-neutralized it shows sensitivity similar to that the original input virus (1, 24–27). Genetic heterogeneity can, however, confer relative resistance to entry inhibition (28). Aggregation of virions with or without antibodies, dissociation of NAbs, failure of NAbs to induce sufficient conformational changes, competition by host-cell receptors, ancillary membrane wrapping of some enveloped viruses, and the degree of cytoplasmic Fc interactions for some naked viruses have all been invoked and partly supported as explanations of the PF. But no general cause is known (1, 23–25, 27, 29–35).

Incomplete neutralization and low slopes of the neutralization curves have been described for various isolates of HIV-1 and simian-human immunodeficiency virus in combination with bNAbs directed to several epitope clusters located all over the envelope glycoprotein trimer, Env: at the trimer apex, the V3-base, the outer-domain mannose patch, the CD4-binding site (CD4- bs), the fusion peptide (FP) and interface between the outer (gp120) and transmembrane (gp41) subunits, and the membrane-proximate external region of gp41 (36–48). Conformational flexibility has been identified as a contributor to PFs in neutralization directed to the trimer apex and to the interface-FP, although such effects may be the atypical features of Env derived from particular isolates (36, 40). In contrast, the exceptionally dense glycan shield on HIV-1 Env is quite plausibly the dominant contributor to epigenetic antigenic heterogeneity, the sources of glycan diversity being variation in occupancy of potential N-linked glycosylation sites (PNGSs) and differential glycan processing (49–61). Indeed, glycan modifications by PNGS mutagenesis, use of expression cells with defects in glycan-processing enzymes, and inhibitors of such processing have implicated glycans in modulating both the potency and efficacy of neutralization (37, 38, 42, 44, 45). In detailed studies of the plateauing of maximum neutralization, the structure of bNAb PGT135 liganded to its epitope on gp120 was combined with site-specific glycan analysis of gp120 to dissect how glycan heterogeneity can lead to bNAb resistance (42, 44). Here, we sought to apply both global and site-specific glycan analyses to explain varying efficacy of neutralization by antibodies directed to other epitopes than that for PGT135. We explored whether virions harboring Env spikes with distinct antigenicities and the corresponding soluble trimer molecules could be segregated by, respectively, affinity depletion and purification and whether neutralization, binding, and glycan analyses could corroborate heterogeneities.

As a model for study, we chose Env of the genotype CZA97.012 of Clade C, the globally most widespread subtype of HIV-1, which is particularly prevalent in Africa and Asia (62, 63). The clone CZA97.012, also known as 97ZA.012, was isolated from peripheral blood mononuclear cells of a 29-year-old female in South Africa and is classified as relatively neutralization-resistant, *i.e.*, of Tier-2 (62, 64, 65). Native-like, soluble CZA97.012 trimer of the SOSIP design can be affinity-purified by bNAbs to quaternary-structural epitopes, including PGT151 to an FP-interface epitope and PGT145 to an apical epitope; purification with the latter, however, results in low yields (39, 66–68).

The PGT151-purified CZA97.012 SOSIP.664 trimer elicits autologous Tier-2 NAbs in rabbits, both in homologous prime-boosting and heterologous boosting (69, 70). The predominant epitope for the autologous NAbs comprises the lining of a hole in the glycan shield around residue D411 in the V4-β19 transition, as shown previously for the heterologous-boost sera (69). Inspection of the autologous neutralization curves for CZA97.012 pseudovirus (PV) revealed variable and sometimes substantial PFs, of up to 60%. We also detected a PF of 15% in neutralization by PGT151 but none with PGT145 or 3BNC117, the latter directed to an epitope mainly overlapping the CD4bs on one protomer but also including the base of the V3 region on an adjacent one (68, 71). Our analyses of CZA97.012 neutralization thus showed a spectrum of drastically different efficacies.

Hence, CZA97.012 PV and the corresponding SOSIP.664 trimer are well suited for testing glycan-based explanations for the variable PF. We confirmed that virions in PV populations have differential reactivities with specific bNAbs by partially depleting PV with PGT145- or PGT151- conjugated beads. We also compared NAb binding to the CZA97.012 SOSIP.664 trimer (66) affinity-purified by different bNAbs. We then explored differences in global and site-specific glycan content in the population of differentially bNAb-reactive trimer molecules by chromatography and mass spectrometry (50–52, 55, 60, 61, 72). The glycan effects were interpreted in their oligomeric- protein context through a novel cryo-EM structure of the CZA92.012 SOSIP.664 trimer in complex with 3BNC117 (68, 71). The ensemble of analyses allowed us to dissect the causes of the variation in PF size.

## RESULTS

### The persistent fraction in CZA97.012 PV neutralization

Neutralization of CZA97.012 PV by PGT145 and PGT151 is shown in **Figure 1**. The results illustrate the disconnect of neutralization efficiency or potency from efficacy or extent. PGT151 was 51-fold more potent than PGT145 at 50% neutralization (IC_50_ values: 0.15 and 7.6 μg/ml, respectively), but less effective than PGT145, leaving a PF at 50 μg/ml of 15%, whereas PGT145 neutralization asymptotically approached 100%. We then combined the two bNAbs at a ratio of 99 to 1, *i.e.,* close to their IC_50_ ratio. Under those conditions, which optimize detection of synergy in potency, only weak synergy was detected at 50% neutralization (combination index, CI = 0.79, Loewe analysis (28, 73, 74)). But at a bNAb ratio of 1 to 1, the combination was weakly antagonistic, possibly reflecting the unidirectional inhibition of PGT151 binding by PGT145 (75) (CI = 1.2). The curves for the individual bNAbs cross at ∼80%, yielding a an IC_80_ ratio of ∼1, and thereby optimal sensitivity for detecting synergy for the combination of 1 to 1 at that level of neutralization. Indeed, at 80% neutralization the synergy was stronger for the ratio of 1 to 1 (CI = 0.52) than 99 to 1 (CI = 0.63). The main observation, thus, is that the synergy was stronger in the zone close to the maximum efficacy of PGT151, a reflection of the visible reduction of the PF (**Figure 1A**).

**Figure 1.**
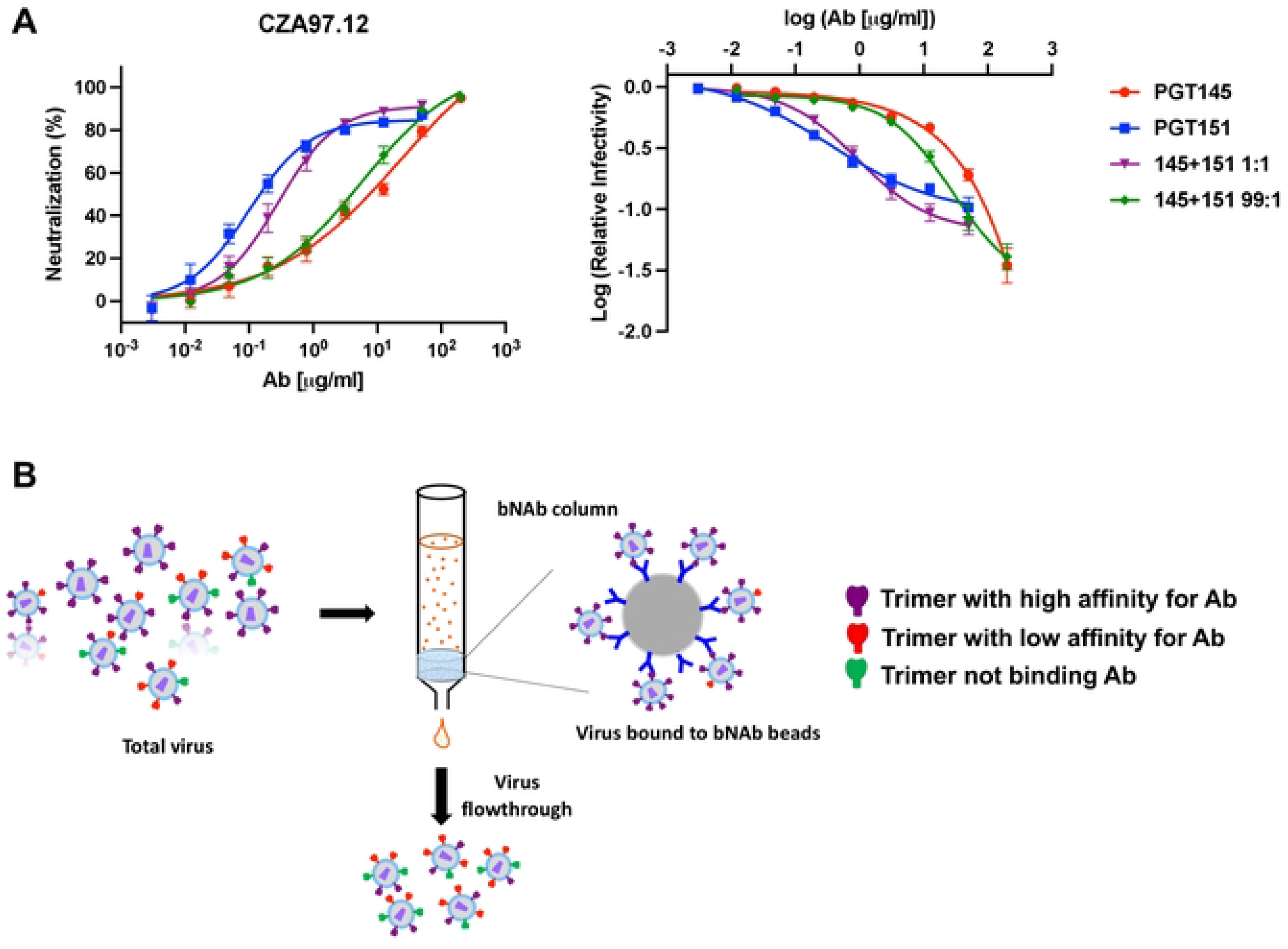
Neutralization and depletion of CZA97.012 PV. A. Extent of neutralization of un-depleted PV (%) in a TZM-bl assay is plotted as a function of the concentration (μg/ml) of PGT145 or PGT151 or their combined total concentration at a mixture of 1 + 1 or 99 + 1 (left). The same results are plotted as the log of infectivity relative to absence of bNAb as a function of the log of bNAb concentration, a plot that better reveals small differences in efficacy and PF (right). The data points in both diagrams are means of 13 replicates ± s.e.m. **B.** The schematic shows preferential depletion of PV particles decorated with Env that binds with high affinity to bNAbs immobilized on Sepharose beads. The unbound virions were then tested in the TZM-bl-based neutralization assay against bNAbs and sera.

A classic Bliss analysis of synergy in extent was not possible at the maxima because of the degree of inhibition by the individual bNAbs was too high (74, 76). Nor can the efficacy of the combination be rationally predicted, because having more than one bNAb molecule bound per trimer may be either redundant or incrementally augment neutralization (77). We therefore simply compared the average residual infectivities (a more general term than PF since it does not require a plateau) for PGT145 and PGT151 with those of the combinations at 50 μg/ml (individual or total bNAb): they were reduced 2-fold at either ratio. The PF was also 2-fold smaller for the combination at a ratio of 1 to 1 than for PGT151 alone and undetectable for the combination at a ratio of 99 to 1, as for PGT145 alone. Hence, enhanced extent of neutralization through combination of the bNAbs was demonstrated.

A plausible explanation both for synergy in potency and for enhanced efficacy would be antigenic heterogeneity, the mechanism being complementary preferences for binding to distinct antigenic forms in the epitope population distributed over the virions (28). We first sought to explore this explanation by differential depletion of PV, as schematically outlined in **Figure 1B**.

### Effects on neutralization of differential depletion of PV by bNAbs

Before performing depletions, we ruled out one potential contributor to the PF: excess of Env in high-dose inocula might absorb NAbs and thereby cause incomplete neutralization. In the case of PGT145 and PGT151 only native-like Env would be relevant. If instead the amount of antigen is negligible in relation to the neutralizing NAb concentrations, the proportion of virus that is neutralized will remain approximately constant over a range of PV-input doses. **SI Figure 1** shows that the drop in log_10_ relative infectivity did not, in fact, vary significantly for neutralization by PGT145 or PGT151 at a constant concentration over a 2.5-log_10_ range of inoculum dose. This result refutes absorption of bNAb by Env as a cause of the PF: it demonstrates that the relative neutralization in this concentration interval adhered to the percentage law of neutralization (78).

The results of the PV depletions are shown in **Figures 2** and **3**. PGT145-depleted PV was highly resistant to PGT145. Conversely, PGT151-depleted PV had drastically enhanced sensitivity to PGT145, the mock-depleted PV falling in between (**Figure 2**).

**Figure 2.**
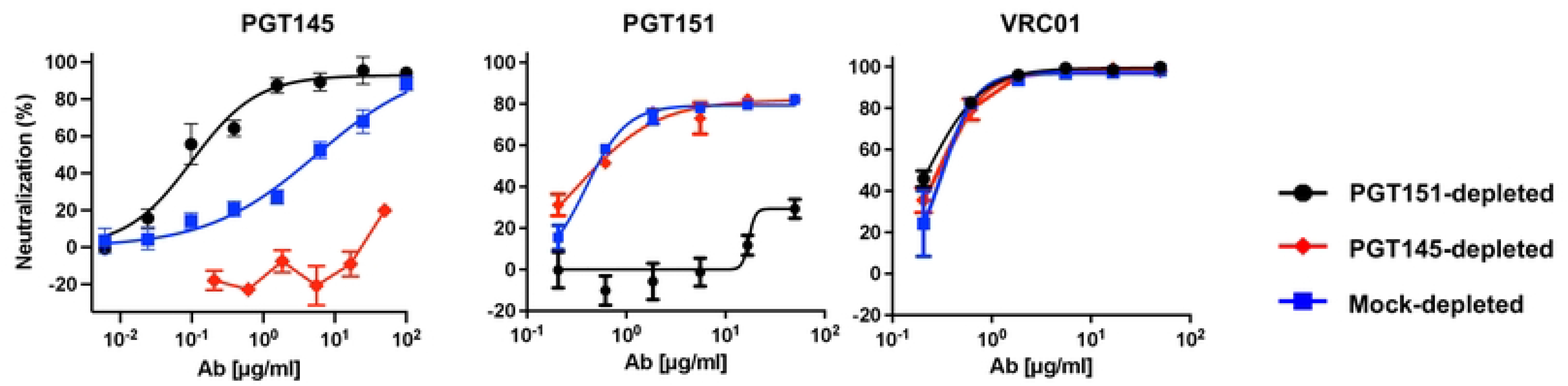
Neutralization of bNAb-depleted CZA97.012 PV by bNAbs. Extent of neutralization (%) is plotted as a function of NAb concentration (μg/ml). PV was first incubated with Sepharose bead-conjugated PGT145 or PGT151 or mock-depleted with control beads (color-coded legend). Unbound virions were then tested against the indicated NAb or serum in a TZM-bl neutralization assay. The data points in the diagrams are means of 2-4 replicates ± s.e.m.

**Figure 3.**
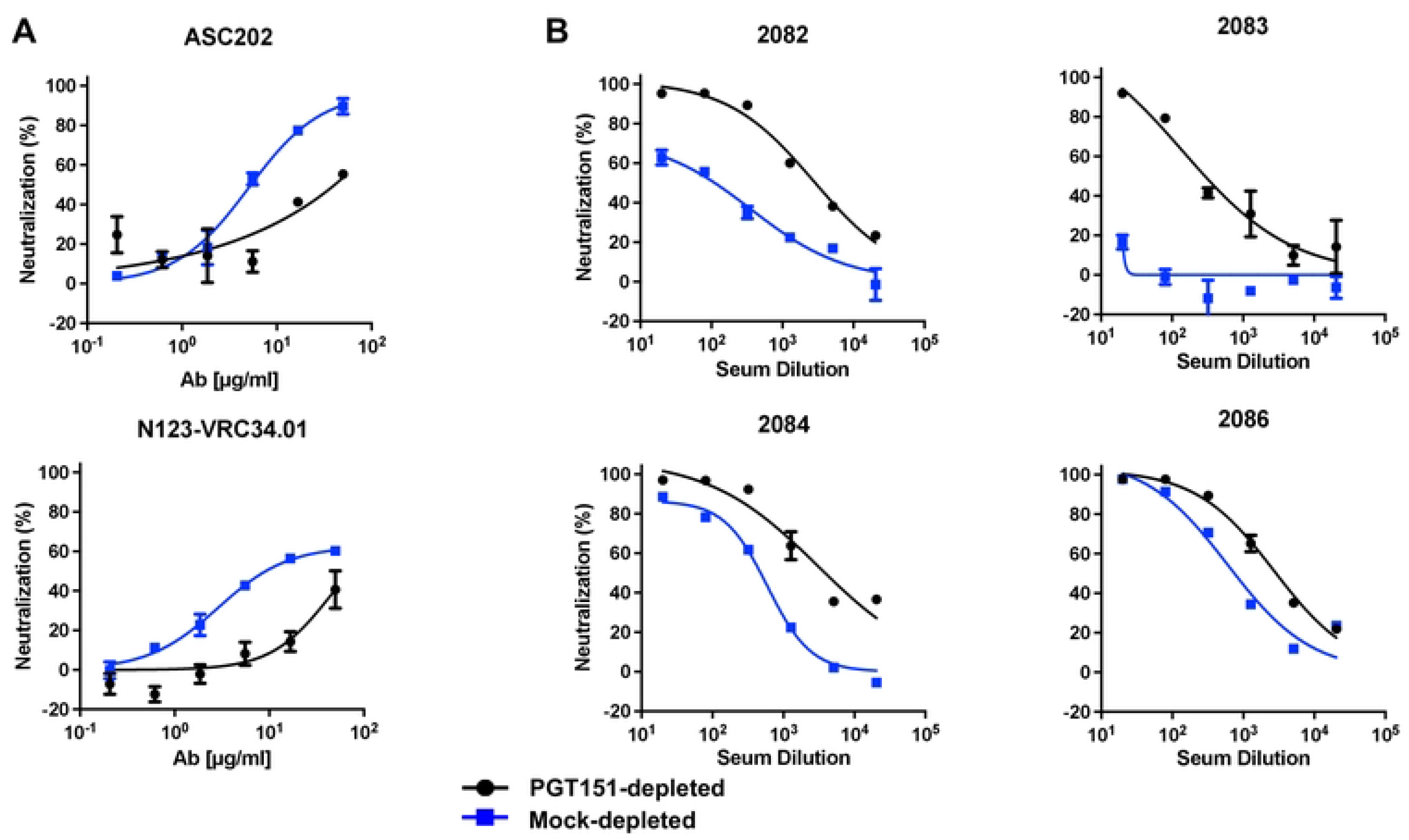
Neutralization of PGT151-depleted CZA97.012 PV by interface-specific bNAbs and post-immunization rabbit sera. PVs were PGT151- or mock-depleted (color-coded legend). Unbound virions were then tested against the indicated bNAb or serum in a TZM-bl assay. **A.** Extent of neutralization (%) is plotted as a function of bNAb concentration (μg/ml). **B.** Extent of neutralization (%) is plotted as a function of x-fold serum dilution. PVs were PGT151- or mock-depleted (color-coded legend). Unbound virions were then tested against the indicated bNAb or serum in a TZM-bl assay. The data points in the diagrams are means of 2 replicates ± s.e.m.

The potency and efficacy of PGT151 neutralization of PGT151-depleted PV was drastically reduced compared with mock-depleted. PGT145-depleted PV, however, differed little in PGT151 sensitivity from the mock control: the potency of PGT151 against PGT145-depleted PV was only modestly greater in the low concentration range (**Figure 2**). The persistent fraction of PGT151 neutralization was indistinguishable, ∼20%, for PGT145- and mock-depleted PV. In contrast, for PGT151-depleted PV the PF was ∼70% (**Figure 2**).

VRC01, directed to the CD4bs (79) neutralized PGT145-, PGT151, and mock-depleted PV indistinguishably within the experimental variation (**Figure 2**).

The effects of PGT145 and PGT151 depletions were thus partly but not symmetrically reciprocal. The results are compatible with a more widespread antigenic heterogeneity in the PGT151 than the PGT145 epitopes on functional CZA97.012-Env spikes. Indeed, little infectivity remained for neutralization analyses after PGT145 depletion, and therefore only PGT151- and mock-depleted PVs were compared in additional analyses with other antibodies.

PGT151 depletion reduced the sensitivity to two other interface-directed bNAbs, ASC202 (80) and N123-VRC34.01 (81) (**Figure 3A**). But it had the opposite effect of markedly enhancing the sensitivity to sera from rabbits immunized with soluble CZA97.012 SOSIP.664 trimer (70). Notably, one such serum, 2083, was non-neutralizing against mock-depleted PV, but upon PGT151 depletion gained neutralizing capacity, which approached 100% efficacy (**Figure 3B**). Autologous rabbit NAbs in such sera are known to be preferentially directed to an epitope lining a defect in the glycan shield on the CZA97.012-Env trimer around residue D411 at the V4-β19 transition (69). Autologous NAbs in the sera collected after homologous prime and boost with CZA97 SOSIP.664 (70), were here found to be largely directed to the D411 epitope (**SI Figure 3)**, as shown previously only for NAbs induced by heterologous-boosts with CZA97 SOSIP.664. It is notable that the serum which showed the weakest effect of PGT151 depletion, 2086, also showed the weakest effect on neutralization of the PNGS-knock-in (KI) mutation D411N: both the PGT151 depletion and the KI mutation affected mainly the potency but barely the efficacy of neutralization by serum 2086. In contrast, the neutralization by the sera 2082 and 2084 was strongly enhanced by the PGT151 depletion and also strongly reduced by the KI mutation (**Figure 3** and **SI Figure 3)**. In spite of the apparent interconnectedness of some of the epitopes studied, however, the KI mutation did not affect neutralization by PGT145, 2G12, VRC01, PGT151, ACS202, or N123-VRC34.01 (**SI Figure 2**).

In conclusion, heterogeneity of the PGT145 and PGT151 epitopes is largely separable among the functional Env spikes on the virions. Lower sensitivity to PGT151 is reflected in lower sensitivity to other bNAbs directed to the FP and interface but linked to higher sensitivity for autologous NAbs directed to the glycan-hole epitope centered on D411. Plugging the latter by a PNGS-KI mutation reduced the sensitivity to the autologous NAbs but not to the tested bNAbs. Hence, we substantiated only a unidirectional connection from the PGT151 to the D411 epitope.

### Effects of differential affinity purification on the antigenicity of CZA97.012 SOSIP.664 trimer

We investigated whether antigenic differences similar to those among virion-associated trimer molecules could be detected also within the population of recombinant soluble CZA97.012 SOSIP.664 molecules (69, 82, 83). Field-flow fractionation in tandem with multi-angle light scattering (FFF-MALS) revealed only subtle differences in the molar masses and hydrodynamic radii of PGT145-, PGT151-, and 2G12-purified trimer molecules (**Figure 4A**). Any actual mass differences would be due to differential post-translational modifications, mainly in glycosylation, whereas the hydrodynamic radius could be affected by glycosylation, conformation, or aberrant folding. The latter was refuted, because trimer preparations purified by 2G12-, PGT145-, and PGT151-affinity and then size-exclusion chromatography showed indistinguishable 100% native-like structures in negative-stain electron microscopy, NS-EM (**Figure 4B**).

**Figure 4.**
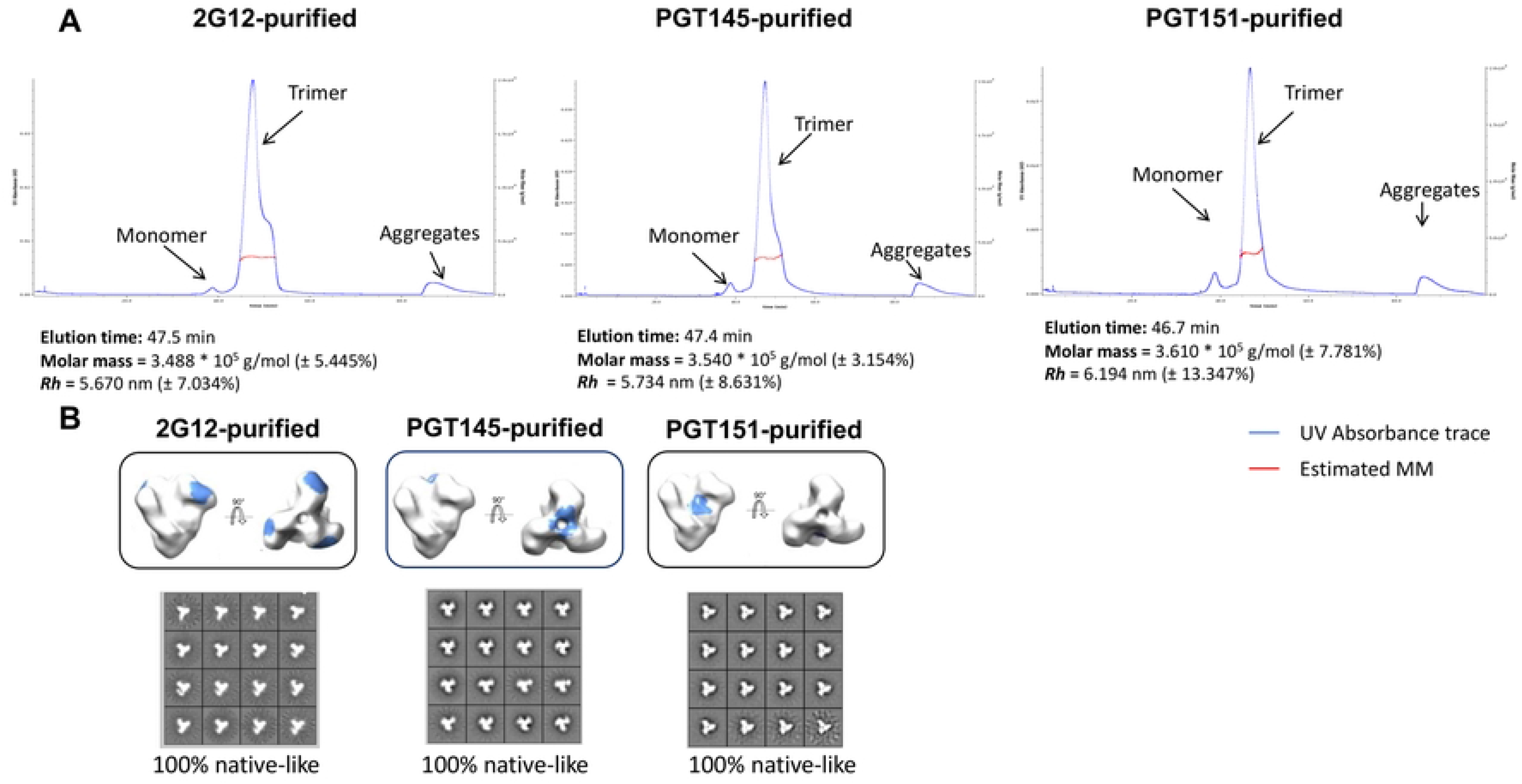
Molar mass and hydrodynamic radius of the CZA97.012 SOSIP.664 trimer measured by FFF-MALS. A. 2G12-, PGT145-, or PGT151- and then SEC-purified trimer was separated by FFF, and the molar mass (g/mol) and hydrodynamic radius (*R_h_*) measured by MALS. **B.** NS-EM was performed on CZA97.012 SOSIP.664 trimer purified with three different bNAbs. CZA97.012 SOSIP.664 trimer was purified by affinity chromatography with 2G12, PGT145, or PGT151 followed by SEC. The epitopes of the three bNAbs are shown in blue on the side and top views of the trimer. 2D-class averages are shown for each purification. The calculated proportions of native-like trimer are given in % below the NS-EM images.

Antigenic differences associated with differential affinity purification of the CZA97.012 SOSIP.664 trimer were first explored by ELISA (**Figure 5**). PGT145 purification gave markedly stronger binding by PGT145 itself than by PGT151 or 2G12, the latter binding being only marginally better than the former. 2G12 purification also gave vastly stronger binding by itself than by the other two, PGT145 binding being marginally better than PGT151. Purification with PGT151 gave strong binding by itself and barely any by PGT145, whereas 2G12 binding was intermediate. The ranking of the trimer binding by two other interface-FP-specific bNAbs, N123-VRC34.01 and ASC202, was the same as for PGT151, but their binding to 2G12- and PGT145-purified trimer was stronger than that by PGT151. Together those results illustrate both similarities and specific preferences among these three bNAbs to overlapping interface-FP epitopes. Thus, **Figure 5** provides strong evidence for stable antigenic heterogeneity in the expressed soluble trimer population: isoforms of the trimer with distinct presentations of each epitope cluster were preserved after elution from each of the purification bNAbs.

**Figure 5.**
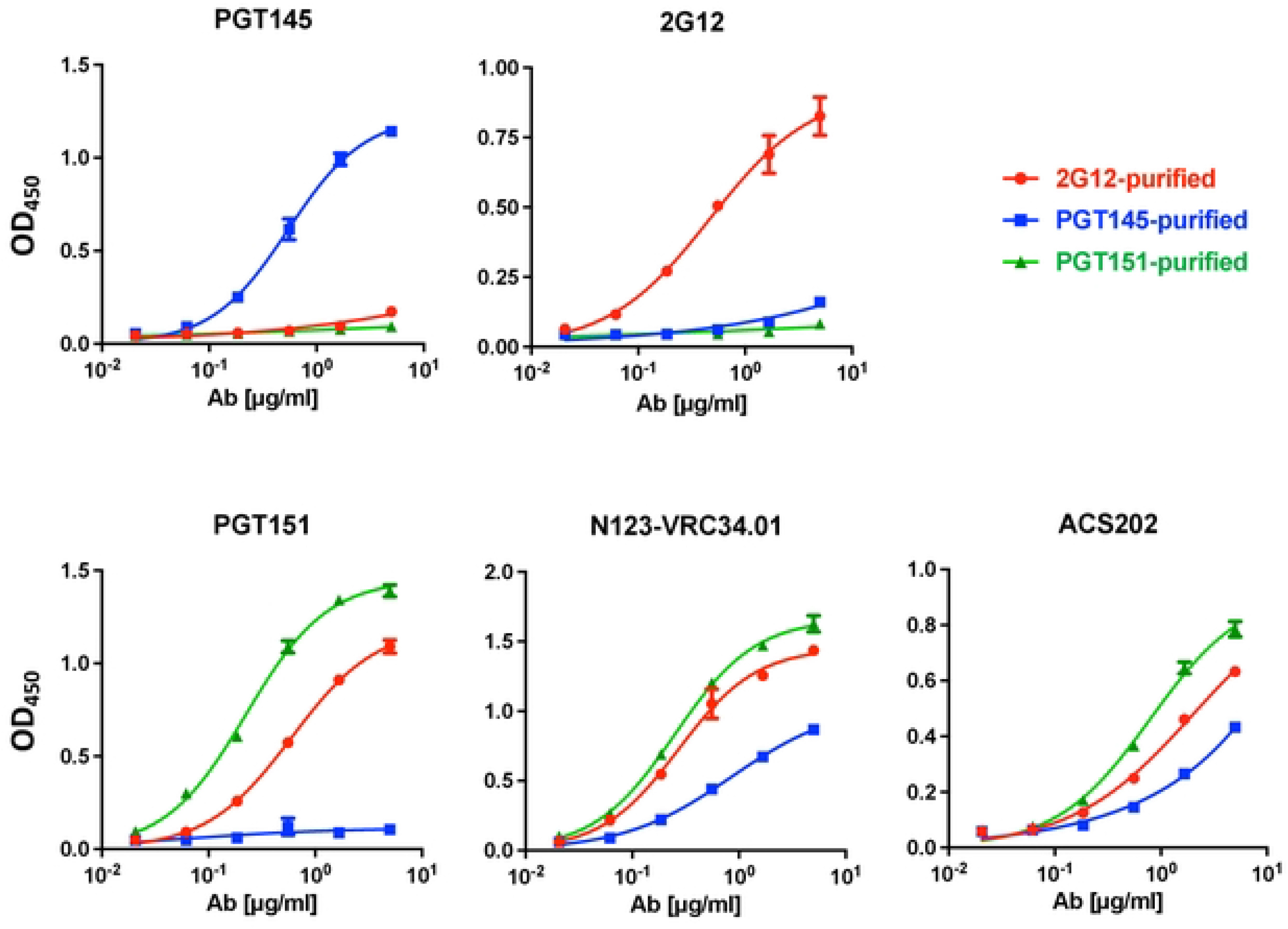
Analysis of bNAb binding to bNAb-fractionated SOSIP trimers by ELISA. Each diagram shows the binding of a bNAb to the CZA97.012 SOSIP.664 trimer purified with three different bNAbs (color-coded legend). The optical densities (OD_450_) after subtraction of zero-bNAb background are depicted on the y axes as functions of the bNAb concentrations (μg/ml). The means of 2 replicates ± s.e.m. are shown.

The antigenicity of differentially purified trimer was then further analyzed by SPR, which unlike ELISA reveals kinetic differences in binding and obviates exposure of some non-NAb epitopes under the ELISA conditions (47, 83) (**Figure 6**). From apex to base, antigenic differences were detected, not just with the purification antibodies. PGT145 bound similarly to PGT145- and 2G12-purified trimer in the association phase but with markedly slower dissociation from the former than the latter. PGT145 binding to PGT151-purified trimer was reduced in both the association and dissociation phase compared with the other two purifications. The faster dissociation of PGT145 from the 2G12- and PGT151-purified trimer may explain the even lower corresponding binding by PGT145 in ELISA, in which the washes amplify loss of antibody through rapid dissociation (**Figure 5**). PG16, also directed to an apical trimer-specific epitope, differentiated an even more clearly: strongest binding to PGT145-purifed, intermediate to 2G12- purified, and weakest to PGT151-purified trimer.

**Figure 6.**
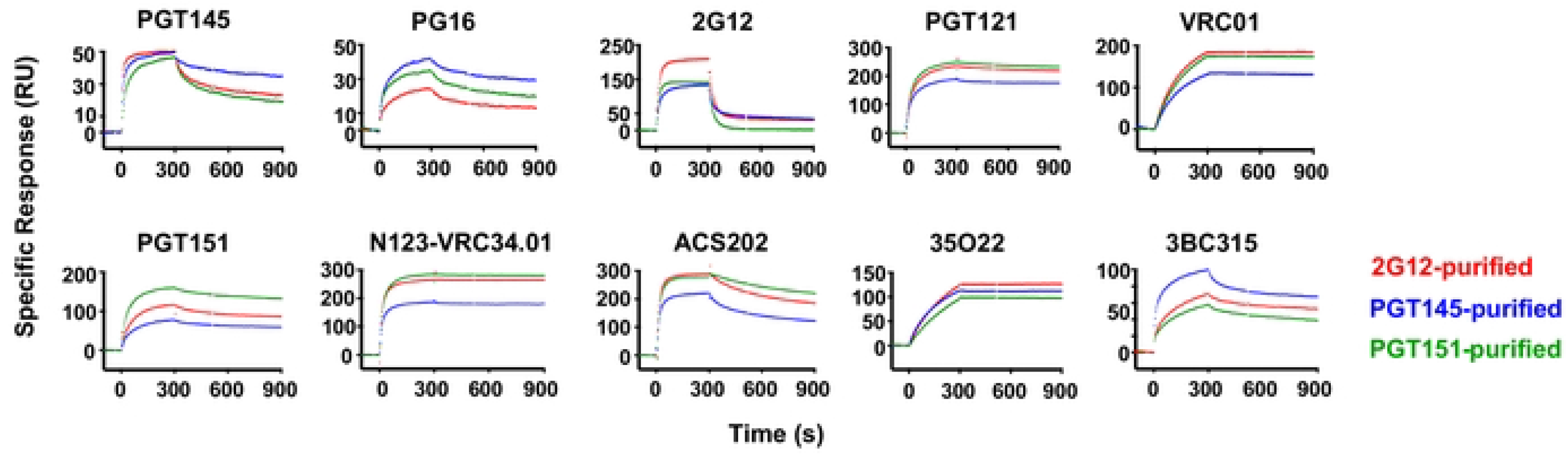
SPR analysis of bNAb binding to bNAb-fractionated CZA97.012 SOSIP.664 trimer. CZA97.012 SOSIP.664 trimer was purified on 2G12-, PGT145-, or PGT151-affinity columns and thereafter by SEC. Sensorgrams show binding to immobilized trimer (response units, RU) after background subtraction on the y-axis *vs.* time (s) on the x-axis. Association was monitored for 300 s, dissociation for 600 s.

2G12 gave highest binding to 2G12-purified trimer, whereas its weaker binding to PG145- and PGT151-purified trimer, differed kinetically, association with and dissociation from the latter being faster (**Figure 6**). The rapid dissociation kinetics of 2G12 may explain its incapacity to neutralize the virus (**SI Figure 2**). Like some other HIV-1 isolates of Clade C, CZA97.012 has both the N332 and N339 PNGSs but lacks the N295 and N392 ones. The net result is the unfavorable kinetics of binding and negligible neutralization by 2G12, whose binding depends on oligo-mannose glycans at these and other sites to various degrees (**SI Figure 2** (66, 84, 85)). Although bivalency of IgG binding contributes little to HIV-1 neutralization strength because of the low Env spike density (86, 87), we note that 2G12 cannot compensate at all for its high off-rate constant by ligating two Env spikes, because its domain swap creates functional monovalency (88). Conversely, when 2G12 is immobilized onto Sepharose beads for affinity purification, the conditions for multivalent interactions do arise: up to 3 epitopes per trimer molecule may ligate 2G12 paratopes. This difference in valency explains why 2G12-affinity columns can be used for purifying CZA97.012 SOSIP.664 trimer, although it does not neutralize the corresponding PV.

PGT121 dissociated extremely slowly from all three trimer preparations but bound at a lower level to the PGT145-purified trimer than to the other two. Thus, the differentially antigenic forms of epitopes at the outer-domain mannose-patch and at the V3-base were selected by the three purification bNAbs, even though two of them engage epitopes elsewhere on the trimer.

As another example of an allosteric effect, the CD4bs-specific bNAb VRC01 showed reduced binding to PGT145-purified trimer, which may be related to the observation that a mutant of CZA97.012 Env shows enhanced PGT145 and reduced VRC01 binding (82).

PGT151 binding gave clearly segregated curves, highest for trimer purified with PGT151 itself, intermediate for 2G12-purified trimer, and lowest for the PGT145-purified one. Two other interface-FP bNAbs, N123-VRC34.01 and ACS202, showed reduced binding to PGT145-purified trimer. The ASC202-binding curves for the PGT151- and 2G12-purified trimers overlapped in the association phase, but ACS202 dissociated faster from 2G12- and PGT145- than PGT151-purifed trimer. A fourth interface bNAb, 35O22, which unlike the others does not make contact with the FP (48, 67, 80, 81), differentiated less but gave yet another ranking: highest binding to 2G12- purified, intermediate to PGT145-purified, and lowest to PGT151-purified trimer. Finally, 3BC315, directed to an inter-protomeric gp41 epitope showed strongest preference for PGT145-purified, intermediate for 2G12-purified, and lowest for PGT151-purified trimer, *i.e.*, the converse of the ranking for PGT151 binding.

In summary, antigenic differences shown in **Figure 6** comprised effects on extent of binding, as well as on kinetics and thereby affinity. They were congruent in that 2G12, PGT145, and PGT151 bound best to the trimer purified with itself. Other differences were registered for epitopes that overlap those for the purification bNAbs. Yet other effects were allosteric, affecting non-overlapping epitopes. Glycosylation differences may be explanatory factors in all these cases, although other covalent modifications cannot be excluded; nor can non-covalent variations on the basis solely of these data (cf. (36)). Allosteric modulation of the antigenicity can arise through domino effects within the glycan shield through varied glycan bulk, charge, and occupancy. Differences in glycosylation may also indirectly affect side-chain and backbone angles (89, 90). We therefore next performed detailed glycan analyses.

### Global and site-specific glycan content of differentially affinity-purified CZA97.012 SOSIP.664 trimer

Variation in glycosylation has been implicated in modulating both the neutralization potency and efficacy of HIV-1 (37, 38, 42, 44, 45). We therefore chose to explore glycan heterogeneity as an explanation specifically of the large PF we observed in neutralization of CZA97.012 by PGT151. Since glycans on Env make direct contacts with all three bNAbs used for purifying the trimers and for depleting the PV, we hypothesized that distinct glycan compositions can be detected after the respective purifications. In addition, we postulated that differentially purified trimer could show distinct glycosylation at other sites that are not in contact with the paratopes, through allosteric effects.

Atypical N-linked glycan processing is abundant on HIV-1 Env. Specifically, the levels of under-processed oligomannose-type glycans are elevated on mature Env (50, 52, 54). Glycan- crowded patches tend to be under-processed and hence contain fewer glycans of the complex type. These under-processed glycans form key components in epitopes for PGT145 and 2G12 (50, 52, 54). The PGT151 epitope, in contrast, contains complex glycans (39, 67, 91). We therefore analyzed N-glycan occupancy and processing on the CZA97.012 SOSIP.664 trimer affinity-purified with each of the three bNAbs.

The trimer was produced in a stably CZA97.012 SOSIP.664-transfected CHO-cell line, which gives high yields and quality (82). To investigate global and site-specific differences in glycan processing among the differentially affinity-purified trimer preparations, we used hydrophilic-interaction ultra-performance liquid chromatography (HILIC-UPLC) and liquid-chromatography mass spectrometry (LC-MS).

We employed a previously reported LC-MS-based method that determines the proportions of oligomannose- and hybrid-type glycans together as one category, complex-type glycans, and N-linked-glycan sequons that remain unoccupied by glycan (**Figure 7** (72)). Oligomannose-type glycans is the collective term for the Man_9_GlcNAc_2_, Man_8_GlcNAc_2_, Man_7_GlcNAc_2_, Man_6_GlcNAc_2_, and Man_5_GlcNAc_2_ moieties (where GlcNAc_2_ denotes *N*-Acetylglucosamine). This analysis revealed an abundance of oligo-mannose- and hybrid-type glycans across gp120 and gp41 in all three purified trimer preparations, most prominently in the intrinsic mannose patch, which in the case of CZA97.012 encompasses glycans at N262, N332, N339, and N386 (54). Furthermore, two of the key V2-apex glycan sites, N156 and N160, were also occupied almost exclusively by oligomannose- and hybrid-type glycans after all three affinity purifications. Although the variation among the differentially purified trimer populations was not drastic, the detected site-specific differences could contribute to the observed differential bNAb binding. The PGT151 epitope comprises glycans at N611 and N637, and micro-array analysis has demonstrated that PGT151 binding favors complex-type glycans (39, 67). In line with that preference, N611 on the PGT151- purified trimer was occupied by <5% oligomannose- and hybrid-type glycans, the favored complex-type glycans making up the vast majority. In contrast, on 2G12- and PGT145-purified trimer, the N611 site was largely occupied by oligomannose- and hybrid-type glycans (∼40% and ∼80%, respectively; **Figure 7**).

**Figure 7.**
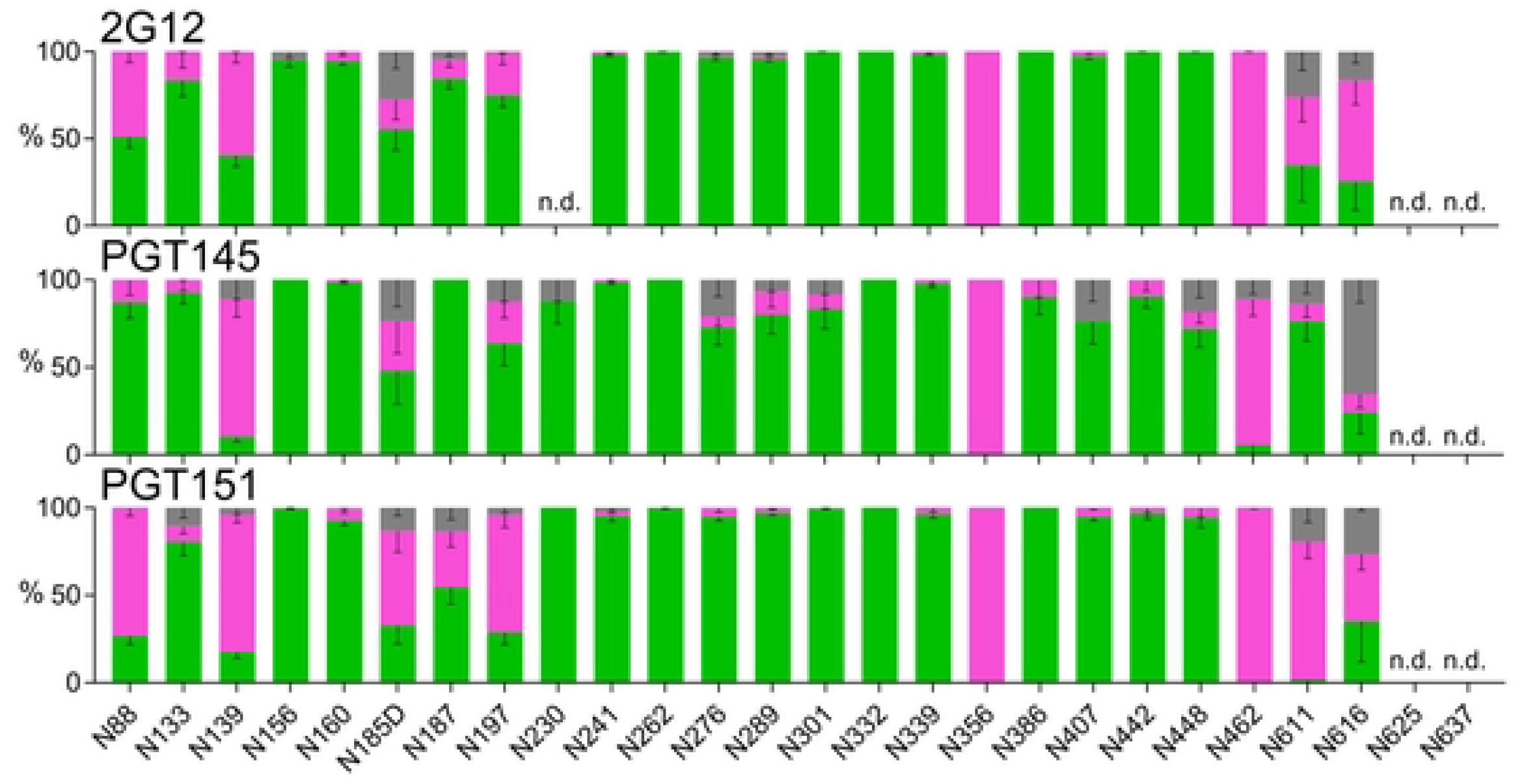
Site-specific glycosylation of differentially affinity-purified CZA97.012 SOSIP.664 trimer. Trimer was purified by 2G12-, PGT145-, or PGT151-affinity chromatography followed by SEC. Differentially purified trimer was analyzed by a method based on PNGase F or Endo H digestion. The bar graphs represent the relative amounts of digested glycopeptides possessing the footprints for oligo-mannose- and hybrid-type glycans together (green), complex-type glycans (magenta), and unoccupied PNGS (gray). The results were obtained by LC-MS. The error bars represent s.e.m. of all peptides detected for each site; n.d. = not detectable (which does not suggest low occupancy).

Comparing the proportions of oligomannose- or hybrid-type and complex glycan and of unoccupancy at each PNGS is informative, but it does not dissect differences in glycan compositions occurring within particular categories. One example of how the subtler subcategories matter is that the outer-domain-directed bNAb PGT135 preferentially binds to oligomannose-type glycans consisting of 8 mannose residues as opposed to 9 (44). To investigate differences in the fine processing of oligomannose-type glycans we first used HILIC-UPLC analysis of the Endo H-released glycan pool, a method which detects global differences in oligomannose-type glycan processing (**Figure 8A**). This approach revealed similar processing of oligomannose-type glycans in 2G12- and PGT151-purified trimer, but significantly higher content of Man_9_GlcNAc_2_ in PGT145-purified trimer. Thus, the observed differences after the three affinity purifications in subsequent bNAb binding quite plausibly result from site-specific fine modifications in glycan processing. We therefore performed further analyses by LC-MS on intact glycopeptides. Because of markedly heterogeneous processing, the coverage of the LC-MS approach was lower, and data could not be obtained for several sites resolved in **Figure 7**. We were, however, able to resolve the key PNGSs of the 2G12, PGT145, and PGT151 epitopes with the exception of N611 (**Figure 8B**).

**Figure 8.**
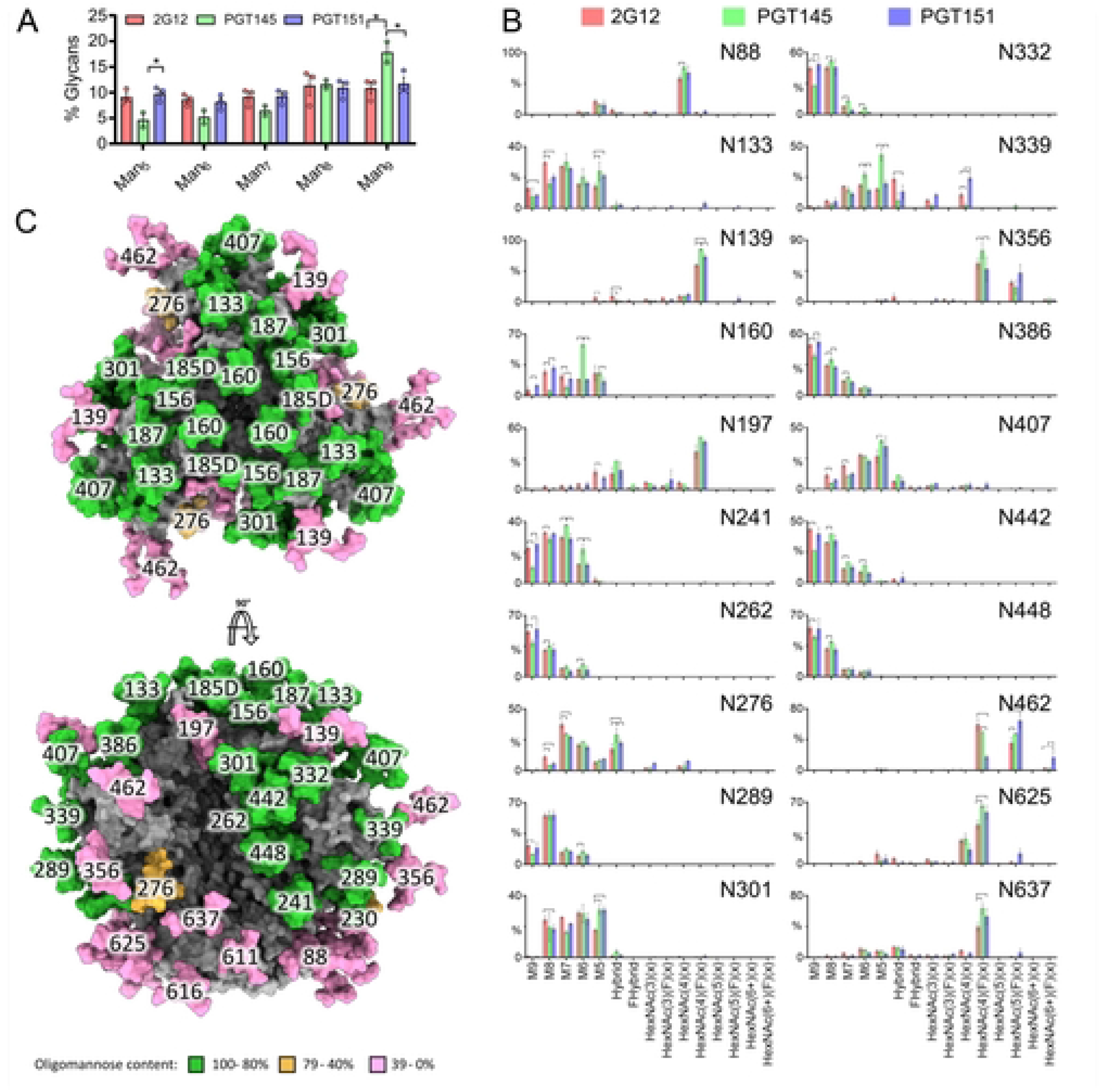
Global and site-specific fine glycan-type determination. A. HILIC-UPLC analysis of Endo H-released and fluorescently labeled N-glycans from CZA97.012 SOSIP.664 trimer purified by 2G12, PGT145, or PGT151 and then by SEC. The bar graphs represent the abundance of each oligomannose-type glycan as listed on the category axis. The percentage on the y axis is proportion of each oligomannose-type glycan out of the total, *i.e.*, PNGase F-released, glycan pool. **B.** CZA97.012 SOSIP.664 trimer was 2G12-, PGT145- or PGT151- and then SEC-purified. Site-specific glycan content was determined by LC-MS on intact glycopeptides. The proportion of each type of glycan is depicted on the y-axes (%), the glycan type on the horizontal category axis. Error bars show s.e.m. for 2 independent MS analyses. Significant differences within pairs of purified trimer preparations (p<0.05, t-test) are bracketed. **C.** Glycosylated model of a HIV-1 Env trimer (truncated C-terminally of residue 664), based on the site-specific glycan data for 2G12- purified material. The predominant glycan composition determined through the analysis displayed in panel B was modelled onto the trimer structure obtained by Cryo-EM of CZA97.012 SOSIP.664 in complex with Fab 3BNC117 and reported in this study (EMD-40088 and PDB 8gje). Where a site was not resolved by LC-MS, a representative glycan consisting of Man_5_GlcNAc_2_ was used instead. Individual glycan sites are coloured according to the abundance of oligomannose-type glycans at each site, as shown in the key.

Man_9_GlcNAc_2_ glycans were elevated at N332 in the 2G12-purfied trimer (**Figure 8B**). The explanation could be that 2G12 makes contacts with α1-2-mannose residues in Man_9_GlcNAc_2_ at N332 and so enriches that glycoform in the trimer population (84, 85, 92). Man_9_GlcNAc_2_ glycans were also elevated in the surrounding glycans (N241, N262, N289, N363, N386, N448) but not at N339, which is a 2G12 epitope component. Since CZA97.012 lacks the PNGSs N295 and N392, which form the primary binding site in the 2G12-BG505 SOSIP.664 complex for which the structure was determined (84, 92), maybe some glycans that are at the periphery of the epitope on BG505 Env instead make direct paratope contact in the CZA97.012 context. That would explain why the preferred Man_9_GlcNAc_2_ glycans were elevated at those PNGSs in addition to N332. But PGT151-purified trimer also had Man_9_GlcNAc_2_ elevations at these sites relative to the trimer purified by PGT145. Hence, an alternative explanation for the differences could be that the Man_9_GlcNAc_2_ glycans in and around the 2G12 epitope perturb the apex. Indeed, at several sites in the intrinsic mannose patch on the outer domain of gp120 PGT145-purified trimer had the lowest content of Man_9_GlcNAc_2_, although overall it had the highest (**Figure 8**). Furthermore, PGT145-purified trimer had significantly higher Man_5_GlcNAc_2_ and Man_6_GlcNAc_2_ content than the other two preparations at N339: the smaller oligomannose-type glycans could favorably affect the positioning of apical residues composing the PGT145 epitope, which overlie the V3 base and adjacent residues. In contrast, complex glycans at N339 were exclusively detected on 2G12- and PGT151-purified forms of trimer (**Figure 8B**).

The most prominent difference occurred at N160, which showed significantly more abundant Man_6_GlcNAc_2_ after purification with PGT145 than with 2G12 or PGT151 (**Figure 8B**). The abundance of Man_6_GlcNAc_2_ agrees with a previous structural study of PGT145 binding to the BG505 SOSIP.664 trimer, which showed that bulky glycans at that site would clash with the antibody (68). Indeed, the density resolved for the N160 glycan in that cryo-EM study agrees with the size of Man_6_GlcNAc_2_, and its core GlcNAc moiety makes contact with the backbone of the descending strand of HCDR3 of the PGT145 Fab; its D1 arm intercalates between HCDR3 and HCDR2, while making polar contacts with the highly conserved residue H52a of the latter region (68).

Notably, the N637 glycan, integral to the PGT151 epitope, showed a significant elevation in glycans containing four N-acetylhexosamines with fucose, which typically corresponds to fucosylated biantennary glycans (HexNac(4)(F)(x)), on both PGT145- and PGT151- compared with 2G12-purified trimer (**Figure 8B**). This type of glycan may favor PGT151 binding but its presence does not explain the antigenic segregation of PGT151- and PGT145-purified trimer populations.

In addition, however, numerous significant differences in precise glycan type among the three differentially purified trimer preparations occurred at PNGSs not intimately involved in any of three bNAb epitopes (**Figure 8B**). These differences can be explained by extensive indirect effects of glycans on antigenicity. They support the ancillary hypothesis, formulated at the beginning of this section, of indirect effects acting across the three purification-bNAb epitopes.

As described above, the sensitivity to the autologous NAbs directed to the epitope around the V4-β19 transition, correlated inversely with PGT151 antigenicity (**Figure 3B**), just as PGT145 and PGT151 antigenicities were inversely related (**Figure 2**). It is therefore noteworthy that the glycans most closely surrounding the V4-β19-glycan defect depended on the purification bNAb in a manner that suggests shielding of the epitope on the PGT151-purified trimer: elevated Man_9_GlcNAc_2_ on N332 and N386, whereas the shorter Man_5_GlcNAc_2_ and Man_6_GlcNAc_2_ on N339 were less abundant than on PGT145-purified trimer, which also had more Man_5_GlcNAc_2_ at N407 than 2G12-purified trimer (**Figure 8B**; **SI Figure 3B**). Hence, these larger oligo-mannose-type glycans would obstruct the autologous-NAb epitope at the bottom of the V4-β19 glycan-hole more on the PGT151- than the PGT145-purified trimer.

CHO cells, although convenient for producing large enough amounts of high-quality trimer for mass spectrometry, differ subtly in glycan processing from HEK-293T cells, in which PV was produced. We therefore also expressed trimer in HEK-293F cells, which give better yields of trimer than do the HEK-293T cells from which they are derived. The glycan composition on CHO- and HEK-293F-cell-expressed trimer was compared by LC-MC. **Figure 9** shows the variation between the differentially expressed forms of trimer in the site-specific oligomannose-type glycan content after the three different purifications, *i.e.*, the sum of Man_9_GlcNAc_2_, Man_8_GlcNAc_2_, Man_7-_ GlcNAc_2_, Man_6_GlcNAc_2_, and Man_5_GlcNAc_2_. Overall, sites displaying an abundance of oligomannose-type glycans did so similarly on the trimer forms derived from the two cell lines, multiple sites containing almost 100% oligomannose-type glycans regardless of cell origin and purification method. That similarity notwithstanding, a modest tendency for higher oligomannose-type glycans on CHO- than on HEK-293F-derived trimer across the purifications occurred; at N133 on PGT151-purified trimer that difference was pronounced. That tendency was also consistent among the three purification methods at N276, where the CHO-derived trimer displayed ∼70% oligomannose-type glycans and HEK-293F-derived trimer only ∼50%. Conversely, we note low oligomannose content at N230, only on PGT145- and PGT151-purifed CHO-derived trimer. Overall, however, the glycosylation of trimer molecules in the two cell lines was similar.

**Figure 9.**
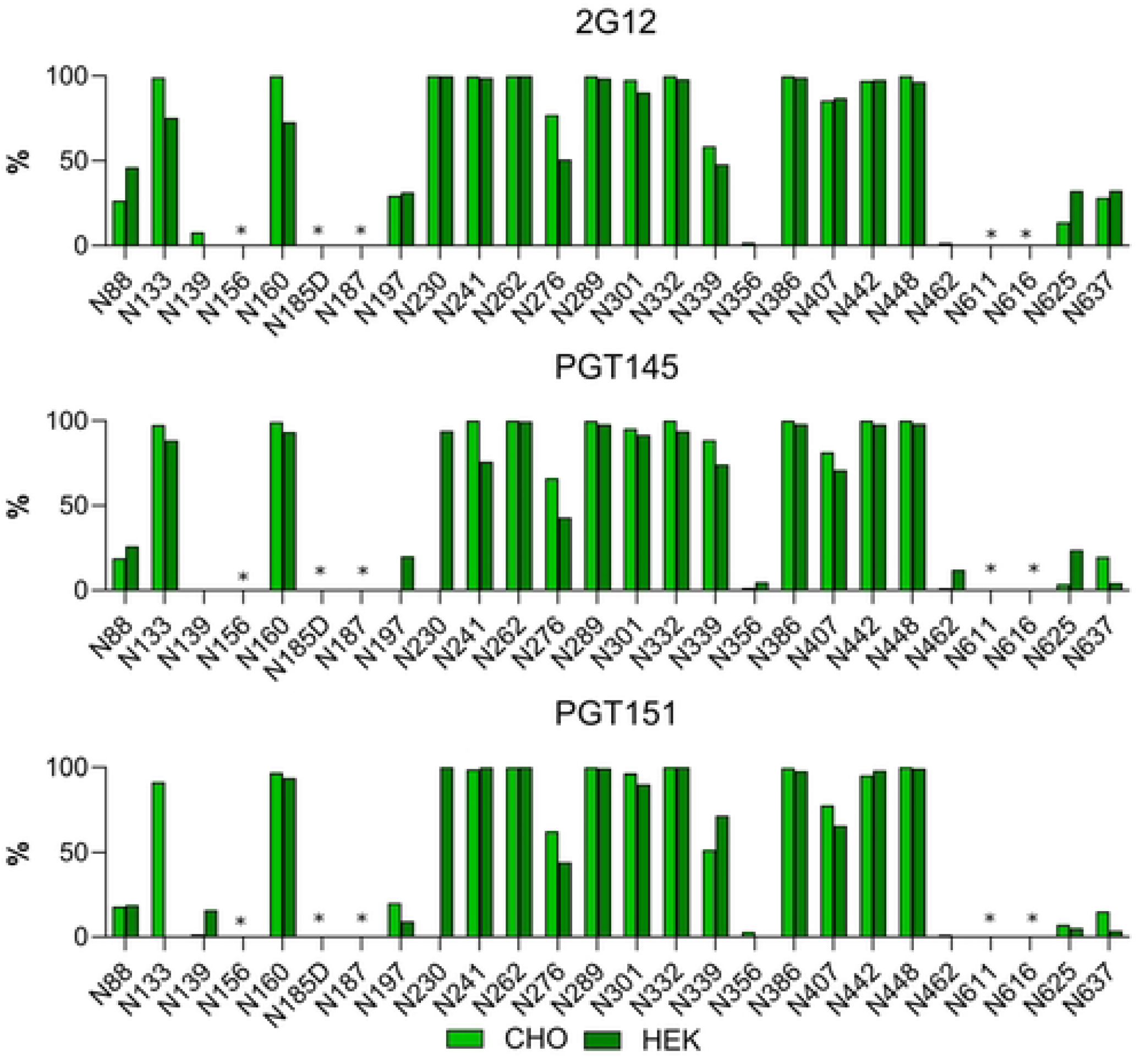
Comparison of CZA97.012 SOSIP.664 trimer produced in different cell lines and differentially affinity-purified. Trimer produced in either a stable CHO cell line or by transient transfection of HEK-293F cells was 2G12-, PGT145-, or PGT151-affinity- and then SEC-purified. Site-specific glycan content was determined by LC-MS on intact glycopeptides as in Figure 8B, but here the contents of oligomannose-type glycans were summed up, *i.e*., only Man_9_GlcNAc_2_, Man_8_GlcNAc_2_, Man_7_GlcNAc_2_, Man_6_GlcNAc_2_, and Man_5_GlcNAc_2_ together, and expressed as percentages of total glycan per site. The bar charts thus represent the relative accumulated oligomannose-type glycan content at each PNGS in CZA97.012 SOSIP.664. Light green represents CHO-derived and dark green HEK-293F-derived trimer (see legend). Sites for which glycopeptides of sufficient quality could not be obtained are left non-determined and are marked with asterisks.

In conclusion, varied glycosylation at sites affecting the 2G12, PGT145, and PGT151 epitopes causes differential affinity purifications by these bNAbs. That segregation is corroborated in binding analyses and explains the differential neutralization potencies and efficacies. Some glycan variations may confer antigenic heterogeneity through direct effects on paratope contacts, whereas others probably influence antigenicity indirectly. The direct and indirect effects may interact in various combinations, together causing substantial antigenic heterogeneity.

### Comparison of 3BNC117 and PGT151 neutralization with the kinetics and stoichiometry of trimer binding by their Fabs

We expanded the analyses to include neutralization by bNAb 3BNC117 and detailed binding properties of 3BNC117 and PGT151. The reason was to complement the comparator PGT145, whose interaction with CZA97.012 Env, on PV and as SOSIP trimer, has some unusual features. PGT145-affinity purification of CZA97.12 SOSIP.664 yields low amounts of trimer (82). Therefore, in a previous study, mutations were introduced into CZA97.12 SOSIP.664 to enhance the interaction, which increased affinity 14-fold. The stoichiometry, however, remained low at 0.48 (82). Since our aim was analysis of binding relevant to neutralization, we did not use those mutations. The moderate affinity of PGT145 Fab for CZA97.12 SOSIP.664 (*K_D_* = 36 nM for a conformational-change model in (82)) agrees with the moderate potency of neutralization (IC_50_ 7.6 μg/ml, *i.e.*, 51 nM IgG in **Figure 1A**). The Hill coefficient of the neutralization curve, however, was low for PGT145 (*h* = 0.45) compared with that for PGT151 (*h =* 0.93, constants derived from data in **Figure 1A**). Since the stoichiometry of PGT145 binding has a maximum of one paratope per trimer, negative cooperativity cannot explain the low slope, which is therefore more plausibly due to antigenic heterogeneity among the virion Env spikes (1, 47, 68, 93). Hence, the Env spikes on virions might have a wide affinity spectrum covering nearly the entire population, whereas ∼50% of the soluble trimer molecules are completely non-antigenic (82), a discrepancy possibly arising from differences in glycosylation between the two trimer populations (61, 72, 94, 95). These features combined render the PGT145-CZA97.012 interaction a complicated outlier. We therefore selected one other bNAb to compare with PGT151, *viz.* 3BNC117 (68, 71). Another reason to study neutralization and binding by 3BNC117 was that its Fab in complex with CZA97.012 SOSIP.664 yielded a high-resolution structure (see below).

The IC_50_ of the neutralization of CZA97.012 PV by 3BNC117 was 0.32 μg/ml; the maximum neutralization was close to 100%. Only in the log-log plot could a tendency towards a PF plateau above 50 μg/ml of bNAb be discerned (**Figure 10A**). Single-cycle SPR analysis showed high affinity, slow dissociation, and a high stoichiometry of 2.8, close to the ideal 3.0 for this bNAb (68); the binding data fitted eminently well to a simple Langmuir model (**Figure 10B and SI Table 1**, single-cycle kinetics; comparison with multi-cycle kinetics, **SI Figure 4 and SI Table 2**). If 3BNC117 IgG bound with the same functional affinity as Fab to Env on PV, the IC_50_ would correspond to an average occupancy of 0.28 on its epitopes. Hence, with random distribution, 63% of the Env spikes would be occupied by at least one bNAb molecule according to the Law of Mass Action combined with binomial analysis (*cf.* (77)). These proportions of virions neutralized and Env spikes occupied correspond to a plausible threshold of neutralization (77). The IC_50_(IgG)/K_D_(Fab) ratio ∼0.40 is compatible with a low but not necessarily negligible degree of bivalent binding. Indeed, IgG neutralizes only marginally or moderately more potently than Fab, with some variation among other HIV-1 PV-bNAb combinations; that the ratios are not higher is attributed to the low Env spike density on virions (47, 86, 87).

**Figure 10.**
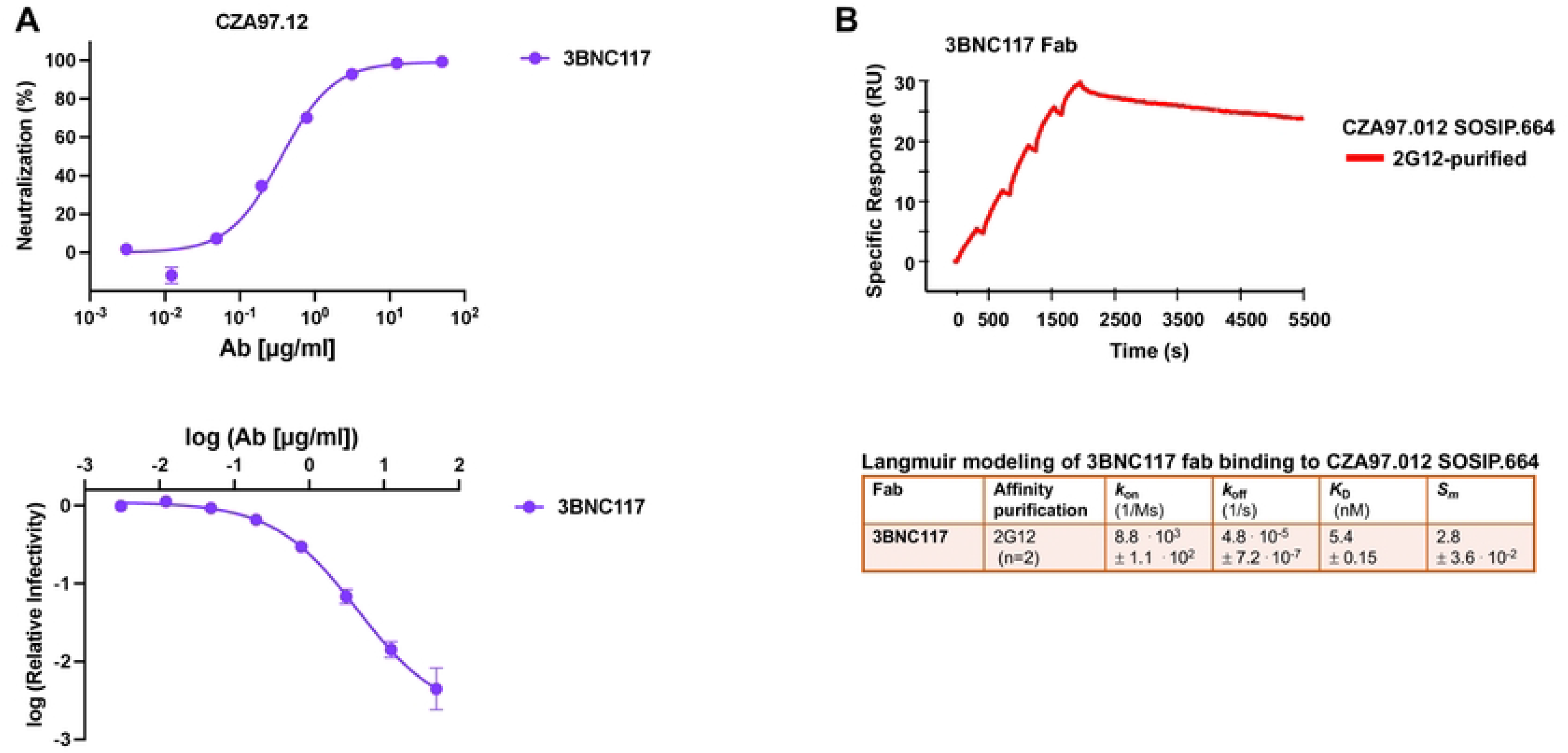
Analyses of neutralization and binding by 3BNC117. A. Neutralization of CZA97.012 PV by 3BNC117 is shown. The diagram depicts a sigmoid curve fitted to % neutralization as a function of bNAb concentration (μg/ml) (top). The log_10_ of the infectivity after bNAb incubation relative to that for absence of bNAb is plotted on the y axis as a sigmoid function of the log_10_ of the bNAb concentration (μg/ml) on the x axis (bottom). The data points in both diagrams are means of two replicates ± s.e.m. **B.** SPR sensorgram for single-cycle kinetics of 3BNC117 Fab binding to 2G12- and SEC-purified CZA97.012 SOSIP.664 trimer. The sensorgram shows the cumulative binding at increasing concentrations to immobilized trimer in response units (RU), after background subtraction, on the y-axis *vs.* time (s) on the x-axis. Each association phase was 300 s; the dissociation at the end was monitored for 3600 s. The number of replicate titrations and the fitted and calculated binding parameters are given in the table underneath the sensorgram.

The binding of titrated PGT151 Fab was also analyzed by SPR, performed as single-cycle kinetics (**Figure 11, SI Table 2**; comparison with multi-cycle kinetics, **SI Figure 4**). Both conformational-change and heterogeneous-ligand models gave considerably better fits than a Langmuir model. To adjudicate between the former two, we performed an injection-time test. The dissociation curves after 30s and 300s association were approximately superimposable, without any tangible retardation of dissociation after the longer association. That suggests an absence of selection for or induction of a better fit (96, 97). We therefore chose the heterogeneous-ligand model (**Figure 7**), which was also validated by the absence of mass-transport limitation and by high T values for all four kinetic constants (**SI Table 1**). Furthermore, the ***k_on1_*** and ***k_on2_*** values differed > 5.7-fold and ***k_off1_*** and ***k_off2_*** values > 35-fold, which means that the kinetics differ substantially between the two sites. The ***K_D1_***and ***K_D2_*** values differed > 210-fold, indicating markedly distinct affinities for the two sites. The lower affinity, however, is not expected to be low enough to generate a sizeable PF, because a site with similar binding kinetics, present as a somewhat larger proportion of the total epitopes, was associated with near-complete neutralization (36).

**Figure 11.**
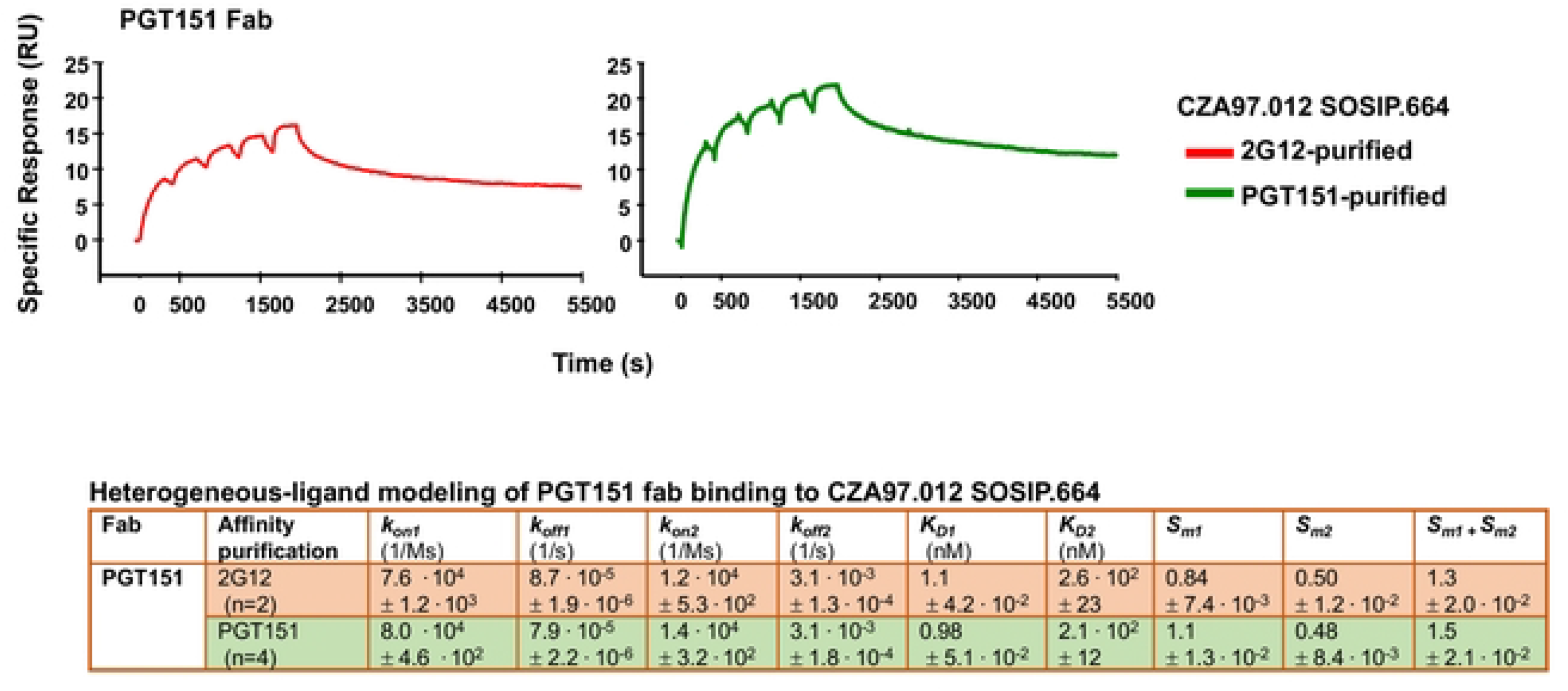
SPR analysis by single-cycle kinetics of PGT151 Fab binding to differentially purified CZA97.012 SOSIP.664 trimer. SPR sensorgram for single-cycle kinetics of 3BNC117 Fab binding to 2G12- or PGT151- and SEC-purified CZA97.012 SOSIP.664 trimer. For comparison, SPR with multi-cycle kinetics, was also performed (**SI** Figure 4). The sensorgrams show the cumulative binding of PGT151 Fab at increasing concentrations to immobilized trimer in RU, after background subtraction, on the y-axis *vs.* time (s) on the x-axis. Each association phase was 300 s; the dissociation at the end was monitored for 3600 s. The number of replicate titrations and the fitted and calculated binding parameters are given in the table underneath the sensorgram.

In contrast to the kinetic constants, the stoichiometric ***S_m1_*** and ***S_m2_*** values differed < 2.3- fold, which means that neither subpopulation is negligible. The cumulative stoichiometry, ***S_m1_*** + ***S_m2_***, was higher for PGT151- than 2G12-purfied trimer (1.5 ± 0.021 *vs.* 1.3 ± 0.020), which indicates that the most drastic heterogeneity in PGT151 antigenicity is reduced by the purification with the same antibody. Why it is not eliminated could have two explanations. First, the avidity difference between the polyvalency in the purification and the monovalency of Fab binding: some interactions with forms of the heterogeneous antigen may be too weak to register by SPR but could contribute to purification through crosslinking of single trimer molecules by two adjacently immobilized copies of PGT151. Secondly, a partial re-equilibration between antigenic conformers may occur after elution. Most important, however, is that for both purifications, stoichiometry of PGT151 binding was substantially lower (*viz.* 1.2 and 1.5) than its ideal value of ∼2.0 measured for SOSIP trimers derived from viruses that are neutralized to near completion, such as BG505 and JR-FL (36, 39, 67).

### Structural interpretation of the antigenic heterogeneity

To put the glycan heterogeneity and antigenicity into structural perspective, we complexed CZA97.012 SOSIP (produced in HEK-293F cells and purified by PGT151-affinity chromatography) with the Fab of the CD4-bs-specific bNAb 3BNC117 (to facilitate orientation sampling) and determined a ∼3.4-Å cryo-EM structure (**Figure 12A** and **B**, (EMD-40088 and PDB 8gje)). Overall, the Env-trimer structure resembles that of previously published ones: the Cα root-mean-square deviation (r.m.s.d.) from BG505 SOSIP.664, also in complex with only 3BNC117, is 1.1 Å (**Figure 12C**). The V4 loop, shorter than in other Env trimers that have been structurally analyzed, is fully resolved, suggesting a more stable antigenic surface, while the aforementioned region around D411 is exposed and flanked by the N332, N339, N386, and N407 glycans (**Figure 12D**). The exposure of the autologous epitope in the glycan defect around D411 explains its high immunogenicity (69). But, in line with differential trimer purification and autologous neutralization results, it would be particularly exposed on the most PGT145-reactive Env spikes, whereas on PGT151-high-affinity trimer molecules, larger oligo-mannose-type glycans would impinge upon it (**Figure 3B, 8B,** and **SI Figure 3**).

**Figure 12.**
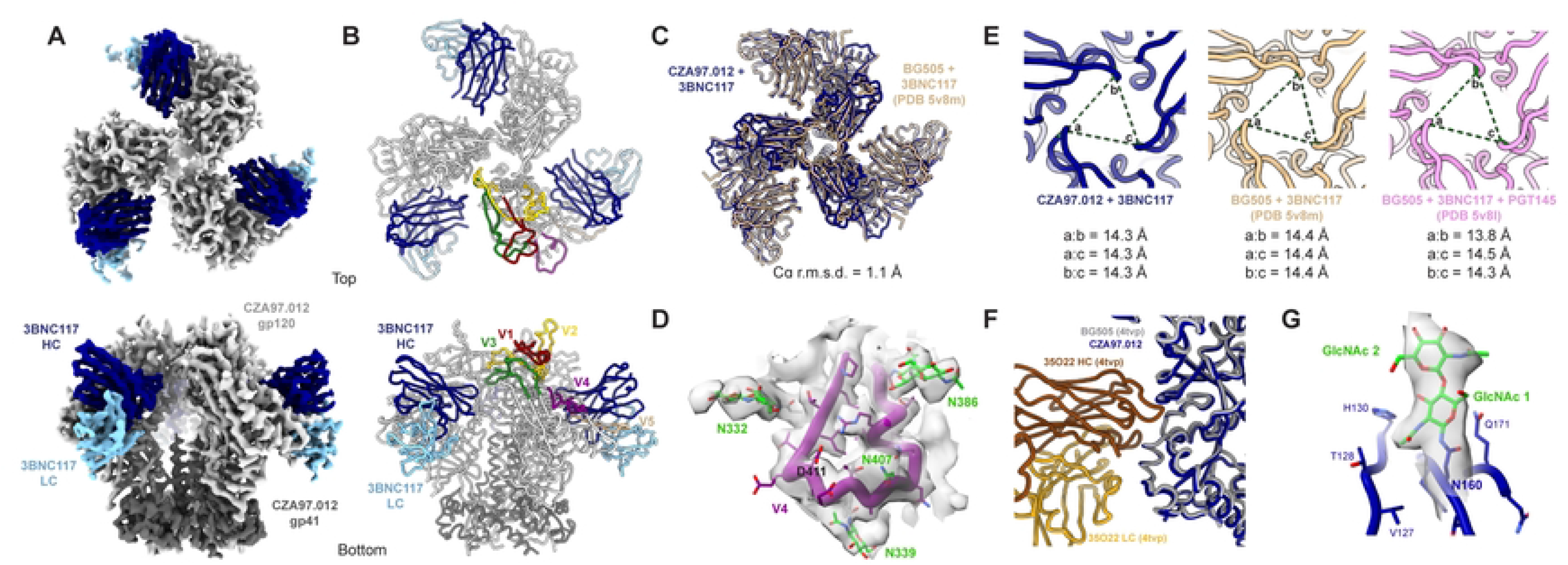
**The CZA97.012 SOSIP.664-trimer structure. A**. The 3.4-Å cryo-EM reconstruction of CZA97.012 SOSIP in complex with 3BNC117 Fab is colored as labeled. **B**. The atomic model of the CZA97.012 complex highlights gp120 variable regions. **C**. Alignment of all 6 Env chains (3- gp120, 3-gp41) of BG505+3BNC117 (PDB 5v8m) onto the CZA97.012+3BNC117 model. Root-mean-square deviation (r.m.s.d.) was calculated between backbone Cα atoms. **D**. Map (contoured at 5σ) and model of the CZA97.012 V4-β19 region and surrounding N-linked glycans are shown. **E**. Interprotomer distances between R166 Cα in CZA97.012+3BNC117, BG505+3BNC117 (PDB 5v8m), and BG505+3BNC117+PGT145 (PDB 5v8l) are given. **F**. Alignment of BG505 SOSIP.664+35O22 (PDB 4tvp (107)) onto CZA97.012+3BNC117, with a focus on the gp41 part of the epitope. **G**. Map (contoured at 5σ) and model of the first two GlcNAc residues of at N160 and surrounding residues of one protomer of CZA97.012 SOSIP.664.

A comparison of the CZA97.012 SOSIP.664+3BNC117 structure with that of the BG505 SOSIP.664+3BNC117 complex and with that of the BG505 SOSIP.664+3BNC117+PGT145 complex (68) demonstrated that the PGT145 epitope was conformationally intact on the CZA97.012 trimer: PGT145-interacting Env residues in BG505 Env are mostly identical in CZA97.012 Env and the opening at the apex – measured as pairwise Cα distances of R166 across protomers – is essentially the same across the 3 structures (maximum difference of 0.5 Å) (**Figure 12E**).

Still, PGT151 purification might somehow affect the dynamics of Env opening and closing through the engagement and subsequent release of the FP. The FP relocates in concert with trimer opening: in the closed state the FP is seemingly solvent-accessible and the N-terminus is disordered in most structures, being intrinsically flexible; in the open state that is induced by sCD4 engagement, the FP becomes more structured and moves towards the core of the trimer (98). Earlier studies showed that pre-binding of sCD4 impedes PGT145 binding: the antibody cannot bind an open trimer (68). If binding and elution from PGT151 perturbed the dynamics of the FP, so that a subset of Env molecules remained more open, then that would decrease binding of PGT145. But cryo-EM of 3BNC117 in complex with PGT151-purified CZA97.012 SOSIP.664 showed only closed trimer, demonstrating that PGT151 purification does not irreversibly open the trimer. The closed CZA97.012 trimer (in the context of bound 3BNC117) presents a similar gp41 topology to that of 35O22-bound BG505 (**Figure 12F**). Furthermore, NS-EM demonstrated that after differential affinity and then SEC purification, CZA97.012 SOSIP.664 was all trimeric (**Figure 4B**). Hence, the observed antigenicity differences do not result from dimer or monomer contamination. Our structural analysis therefore concludes that PGT151-purified CZA97.012 SOSIP.664 resembles native-like trimer in the areas of its bNAb-epitope clusters and maintains a closed conformation.

Density for the first two GlcNAc residues of the N160 glycan is visible in the cryo-EM map, suggesting that PGT151 purification is not selecting for glycan underoccupancy at this key PGT145 site. Any such selection would have had to be allosterically mediated because of the distance from the PGT151 epitope to trimer apex (**Figure 12G**).

Taken together, the observations of a closed conformation and of full glycan occupancy at N160 of PGT151-purified trimer suggest that differences in binding and neutralization after trimer purification or PV depletion with PGT145 and PGT151 therefore most likely stem from the other glycosylation heterogeneities at the respective epitopes that we describe above (**Figures 7-9**).

In summary, variation in the occupancy by and composition of glycans at sites affecting the 2G12, PGT145, and PGT151 epitopes is a plausible contributor to differential binding and neutralization efficacies and potencies. Some glycan variations confer antigenic heterogeneity through direct effects on paratope contacts, while others may do so allosterically.

## DISCUSSION

We found that clonal CZA97.012 Env trimer molecules, both on PVs and as soluble SOSIP protein, contain antigenically distinct forms of the PGT145 and PGT151 epitopes. We infer that the populations of Env-trimer molecules have overlapping spectra of antigenicities with discrete maxima for these bNAbs. Glycan composition differed significantly at 20 PNGSs on CZA97.012 Env, including glycans crucially contributing to the epitopes of the affinity-purification bNAbs. Therefore, both direct and indirect effects of glycan differences on the epitopes can be deduced. How can glycosylation and conformation as causes of the PF be dissected? Here, by FFF-MALS, we observed distinct masses and hydrodynamic radii of soluble trimers affinity-purified with the different bNAbs. Whereas the radii could change along with the conformation, significant mass differences would incontrovertibly demonstrate differences in post-translational modifications, largely comprising variations in glycan-site occupancy and glycan processing (49–54, 56). The differences in mass detected by FFF-MALS among the differentially purified trimer populations were, however, merely compatible with differential processing; they were too small to prove it. But the glycan analyses were definitive on this point. Specifically, at N160, PGT145- purified trimer had less Man_9_GlcNAc_2_, Man_8_GlcNAc_2_, and Man_7_GlcNAc_2_ but more Man_6_GlcNAc_2_ than trimer purified by the other two bNAbs, and more Man_5_GlcNAc_2_ than on PGT151-purified trimer. These observations agree with structural indications of which type of glycan is best tolerated by the PGT145 paratope at that site, because cryo-EM of BG505 SOSIP.664 in complex with PGT145 Fab showed a density at N160 strongly resembling Man_6_GlcNAc_2_ (68).

Most important for explaining the large PF in PGT151 neutralization was the low stoichiometry, detected by SPR, of 1.3 PGT151 Fabs per 2G12-purified trimer (**Figure 11**). This value is substantially lower than the ideal binding of two PGT151 Fabs per trimer observed with versions of the BG505 SOSIP trimer (47, 67, 99). The reduced stoichiometry could arise as an average of different proportions of occupancy by 0 or 2 or Fabs or possibly 1. Even the PGT151- purified trimer did not show the ideal stoichiometry of 2, but an intermediate one of 1.5, which suggests some trimer molecules eluted from the affinity-chromatography column could bind only 1 Fab. The corresponding proportion of un- or underoccupied Env spike on virions could be enough to cause a sizable PF (77).

Which glycan differences within the trimer population could explain the reduced stoichiometry of PGT151 binding? A complex glycan at N611 directly interacts with the paratope (67). BG505, which is neutralized without any detectable PF by PGT151 (36), has exclusively complex glycans at N611 both on membrane-anchored Env spikes and SOSIP.664 trimer (61, 67). Notably, we observed equal proportions of oligomannose- or hybrid-type and complex glycan on CZA97.012 SOSIP.664 at N611 after the non-segregating purification by 2G12, a majority of oligomannose- or hybrid-type glycan after PGT145 purification, and exclusively complex glycan after PGT151 purification (**Figure 7**). The oligomannose- or hybrid-type glycan at N611 in the mixed trimer population quite plausibly reduces the average stoichiometry of PGT151 Fab binding (**Figure 11**), and thereby causes the large PF (**Figure 1**). Full N611 occupancy by particular complex glycans would be conducive to the ideal stoichiometry of two paratopes per trimer (**Figure 1** and **7**). Additionally, the observed heterogeneity in the precise type of complex glycan at N637 among the antigenic forms purified by the respective bNAbs, perhaps in combination with allosteric glycan effects, could modulate stoichiometry and thereby neutralization efficacy (**Figure 8**).

A potential caveat in these dissections of PF causality is that differences in glycosylation between certain sites on soluble and viral-membrane-anchored trimer molecules have been observed for some strains (61, 94). But those are not uniform occurrences. Much evidence supports eminent similarity in antigenicity and glycosylation patterns between membrane anchored Env spikes on virions and several SOSIP trimers studied (47, 48, 61, 80, 83, 94, 98, 100–108). We therefore consider the following explanatory scenario justified. A primordial heterogeneity in glycosylation arises in the endoplasmic reticulum as varied occupancy and in the Golgi as varied type of glycan. This variation would then manifest itself in antigenic heterogeneity through direct paratope contact effects and allosteric effects on glycan and amino-acid side-chain orientation, as well as backbone conformation. Such combined net effects would increase or decrease the stoichiometry of binding. Ultimately, the stoichiometry falls above or below a minimum threshold of paratopes bound per virion required for neutralization (1, 7, 23, 77, 109–112).

It may be fruitful to compare how the PF is caused in PGT151 neutralization of B41 (36) and of CZA97.012. Through structural studies, we could attribute the large B41 PF to the pronounced conformational flexibility of the B41 trimer (83, 98, 113), the opening of the trimer yielding either steric clashes with the PGT151 paratope or the sequestering of a key component of the epitope, *viz.* the FP (36). Although the trimer opening would be a sufficient cause of the large PF, the propensity for a sub-population of trimer molecules to assume or stay in an open conformation can have a number of antecedent causes, including allosteric effects of heterogeneous glycosylation (36). But B41 may be a special case: the conformational flexibility of its Env protein is unusual among the different HIV-1-derived trimers studied so far (98, 113). Large PFs in PGT151 neutralization, in contrast, are common (36, 39). Thus, the intriguing explanation for the large PF in PGT151 neutralization of B41 (36) lacks the requisite generality to explain a fairly frequent phenomenon. Here we present a hypothesis about PGT151 neutralization that does not: our findings support direct and indirect effects of widespread heterogeneity in glycosylation, which stems from the high glycan density on Env and myriad stochastic fates along the secretory pathway. The hypotheses imply discrepancies in global glycan composition, heterogeneity in paratope-contacting glycans, and multiple glycan differences with potential allosteric effects. On all counts we found evidence for such variation, which would give antigenic heterogeneity and thereby to PF variations (**Figures 7** and **8**).

How do the findings about the PF and glycosylation inform immunogen design? Just as for B41, which also shows a large PF in PGT151 neutralization (36), the PGT145 and PGT151 epitopes on the CZA97.012 trimer seem allosterically interconnected. In the case of both virus isolates, their antigenic maxima are discrete in the trimer population, and antigenic variants with lower PGT151 affinity present the main autologous epitope in a more accessible or paratope- fitting form than do the average trimer molecules. But the epitopes for autologous neutralization are located in distinct regions on the B41 and on CZA97.012 trimers. Both of them, however, line defects in the glycan shield: on the B41 trimer around residues 230 and 289 and on the CZA97.012 trimer around residue D411 at the V4-β19 transition (69, 114, 115).

The inverse relationship between PGT151 binding and autologous neutralization was here further strengthened by association between the magnitude of the effect of PV depletion by PGT151 and of the D411N-KI mutation on the neutralizing capacities of the autologous rabbit sera (**Figure 3** and **SI Figure 3**). The interconnectedness of the bNAb and autologous epitopes might suggest allosteric effects of plugging the glycan defect, but we found no evidence for that: the bNAbs tested neutralized the KI mutant and the parental virus similarly (**SI Figure 2**). Nevertheless, if preferential presentations of the autologous and the PGT151 epitopes by isoforms are incompatible, which form would best induce breadth? Efforts to broaden autologous neutralization responses to partly overlapping glycan-defect epitopes on B41 and BG505 Env seem to abut intricate molecular barriers (115). If those responses are not amenable to broadening, they must be considered off-target (116). Do the responses to the glycan-defect epitope centered on D411 on CZA97.012 Env hold any greater prospect for broadening?

The high-resolution structure of the previously uncharacterized CZA97.012 SOSIP.664 trimer we present here may help answer that question (**Figure 12**). In contrast to other structurally analyzed Env trimers, the CZA97.012 trimer has a fully resolved V4 loop, which suggests a relatively stable antigenic surface there, probably because the shortness of this loop, a characteristic of Clade C isolates (117). This greater stability might favor certain broad but clade-restricted NAb responses (118–121). Whether this stability and exposure are conducive to eliciting intra-Clade breadth or merely promote off-target narrow responses cannot be determined by our data. But the structural information may assist the design of mutants that induce broadened autologous responses, without compromising the structural integrity of known cross-Clade bNAb epitopes (cf. (115)).

Since preferential PGT151 binding and neutralization were linked to similarly superior activities of other FP-directed bNAbs (**Figures 3**, **5**, and **6**), induction of such bNAbs might be favored by PGT151-purifed immunogens. Furthermore, PGT151-affinity purification reduced PGT145 binding less than *vice versa*, whereas PGT145-affinity purification uniquely reduced PGT121 and VRC01 binding. Together, these effects suggest a net advantage of PGT151-affinity purification. Conversely, 3BC315 binding to its epitope was better after PGT145 purification. It is desirable to induce bNAbs of many specificities, for reasons we conclude with below. Therefore, mixtures of differentially purified immunogens, or those purified with, *e.g*., 2G12 and SEC (122), which segregates antigenic forms less than do PGT145 and PGT151, might be optimal.

Large PFs *in vivo* would be problematic after both the successful induction of bNAbs by active immunization and the administration of bNAbs as passive immunization for prevention or therapy (10, 12, 123). Several factors may conspire to reduce both potency and efficacy of neutralization *in vivo* compared with *in vitro*: PV is more sensitive to neutralization than fully replicating virus, and virus derived from transfected 293T cells is more sensitive than that derived from primary lymphocytes, which is in turn more sensitive than virus produced in macrophages (124–126). The latter difference has been attributed to N-linked lactoseaminoglycans, present exclusively on macrophage-derived Env (125). In addition, HIV-1 is more genetically diverse *in vivo* than *in vitro*, which raises the bar for entry inhibition (28).The size of the PF after exposure of vaccinees or recipients of bNAbs for prevention could make the difference between sterilizing immunity and viral breakthrough (17, 36, 123).

The PF in therapeutic passive immunization could undermine the suppression of viremia and the elimination of recalcitrant viral reservoirs. The outgrowth virus isolated from the HIV-1 reservoir after bNAb therapy is often largely sensitive to the treatment bNAbs (127–129).

Therefore, a contributing explanation to what maintains the reservoir might be epigenetic resistance, conferring incomplete neutralization by the treatment bNAbs.

Our findings of a substantial synergy in potency of combined bNAbs, specifically in the zone of higher degrees of inhibition (*i.e.,* at *IC_80_*), and of PF reduction by a bNAb combination, together indicate the advantage of having multiple bNAb specificities both in active and passive immunization (130–134). The synergy is explained by partly segregated differential antigenicities over the virion population (1, 28, 77). Such a mechanism can even account for synergy between NAbs directed to the same epitope cluster, provided the trimer population on virions presents antigenic variants that interact differentially with those NAbs (135).

To conclude, the PF in neutralization can be explained on the basis of antigenic heterogeneity, created by multiple post-translational modifications and conformational variability, and reflected in reduced stoichiometry; it can be minimized by combining multiple bNAb specificities.

## MATERIALS AND METHODS

### PV production

HEK-293T cells, maintained in growth medium (DMEM with 10% FBS, 2mM L-Glutamine, and 1% penicillin-streptomycin), were used for producing PV. One day before transfection, cells were seeded in 6-welled plates at a density of 4 × 10^5^ cells/well. Transient transfection was performed with ProFection Mammalian Transfection System (Promega), which is calcium-phosphate-based, in accordance with the manufacturer’s protocol. Three hours before transfection, the medium of 50-60%-confluent cell cultures was changed to antibiotic-free. Cells were transfected with CZA97.012 *env* pcDNA3 and pNL4.1AM *env* (-) *luc* (+) plasmids (ratio of 1:2) diluted in CaCl_2_ (2M) and mixed with transfection reagent in 2 × saline solution buffered with 4-(2-hydroxyethyl)-1- piperazineethanesulfonic acid-buffered (HEPES). The transfection mix was added to the cells and incubated at 37^°^C in air containing 5% CO_2_. The next day, cells were supplemented with fresh growth medium and incubated further for 48h. PV was harvested 72 h post-transfection by spinning the supernatant at 2000 rpm for 10 min. An additional 10% FBS was added to the supernatant, before spinning, to maintain virion integrity.

### TZM-bl neutralization assay

Neutralization of PV by bNAbs and heat-inactivated sera was analyzed in a TZM-bl-cell-based assay, as described (136). The day before infection, cells were seeded in 96-well plate (white tissue-culture treated, Costar 3917), at a density of 1 × 10^4^ cells per well. PV (diluted in DMEM growth medium to yield luminosity readouts of ∼2000 000 counts/s) were incubated with serially diluted bNAbs or sera (heat-inactivated) for 1 h at 37^°^C. The high dose was chosen to obtain a wide dynamic range, which allows sensitive PF identification. We verified complete absence of bleed-through of signal to neighboring wells, even at this high viral dose. The mixtures of PV and bNAb or serum were transferred to TZM-bl-cell cultures. Three days after infection, medium was aspirated and the cells were lysed with Glo-lysis buffer (Promega). Plates were frozen for at least 2 h at -80^°^C. After thawing, Bright-Glo substrate was added to each well. The luciferase signal was read on an Enspire multimode plate reader (Perkin Elmer). Relative infectivity was analyzed by subtracting the background signals (cells but no virus) from signals of virus-cell wells. The neutralization (%) in relation to the infectivity of virus without Ab was calculated for all bNAb and serum dilutions. The signal in the absence of antibody was stipulated to represent 100% infectivity, *i.e.,* 0% neutralization. Data were analyzed and plotted with Graph Pad Prism 6 software; four-parameter sigmoid curves were fitted to the data with maximum constrained to 100 %. The assay described above was used to generate the results in **Figures 1-3**, **10**, and **SI Figures 2-3**. In **SI Figure 1** a variant of the assay was used, with a constant bNAb concentration (50 μg/ml) and a titrated viral input. The D411N mutant (**SI Figures 2** and **3)** has been described previously in a study mapping of autologous neutralizing responses of rabbits to CZA97.012 SOSIP.664 immunization (69).

### Depletion of PV by bNAbs

PV was added to affinity columns containing Sepharose beads with covalently coupled PGT145 or PGT151 or without Ab (mock control). The bNAb affinity columns were made by cross-linking IgG to activated CNBr-Sepharose 4B beads (GE Healthcare) according to the manufacturer’s instructions and as previously described (83). The columns containing PV and beads were incubated at 37^°^C for 3 h with constant mixing on a nutator. Initially, to assess the effect of temperature and nutation, aliquots of PV without beads, designated un-depleted PV, were subjected to the same treatment. Un-depleted and mock-depleted PV later showed indistinguishable neutralization, and the un-depleted control was discontinued. The mix of PV plus beads was allowed to settle at room temperature in vertical columns. PV was collected by gravity flow-through from the respective column, filtered through 0.45-μm membranes, and immediately analyzed in a neutralization assay.

### Expression and selective purification of native Env trimer

First, trimer was produced and differentially purified from a CZA97.012 SOSIP.664-expressing CHO-cell line maintained in ExpiCHO Expression Medium (Thermo Fisher Scientific) (82). Then, the CZA97.012 SOSIP.664 trimer was also expressed transiently in HEK-293F where indicated. Cells were cultured in Expi293 Expression Medium (Thermo Fisher Scientific) supplemented with 1 × penicillin-streptomycin. The construct CZA97.012 SOSIP.664 with His-tag (GSGSGGSGHHHHHHHH) was cloned into the pPPI4 expression vector (GenScript). HEK-293F cells were transiently transfected by the use of 293fectin (Thermo Fisher Scientific). Briefly, cells were seeded at a density of 4 × 10^6^ cells/ml in 250 ml of medium (with antibiotics), a day before transfection. On the day of transfection, cells were suspended in antibiotic-free medium at 6.0 × 10^6^ cells/ml with 90% viability as measured by trypan blue exclusion. *CZA97.012 SOSIP.664* and *furin* plasmids were diluted in Opti-MEM at a ratio of 4:1, mixed with 293fectin reagent, and incubated at room temperature for 20 min. The 293fectin-DNA mix was added to the cells. Transfected cells were incubated at 37°C with 5% CO_2_ and constant shaking for 3 days. Then cells were centrifuged at 3000 rpm for 30 min. Supernatant was collected, passed through a 0.2-μm filter in preparation for 2G12-, PGT145-, and PGT151-affinity purification. bNAb affinity columns were made by cross-linking IgG to activated CNBr-Sepharose 4B beads (GE Healthcare) according to the manufacturer’s instructions. The filtered supernatant was allowed to pass through the affinity columns. Bound trimer was eluted with 3M MgCl_2_, as described elsewhere (82). MgCl_2_ was removed by dialyzing through snake-skin tubing in an exchange buffer (TN150: 20mM Tris-HCl, 150mM NaCl, pH 8) overnight at 4^°^C. SEC with TN150 as running buffer was then performed on a Hi Load 16/600 Superdex 250 pg column, to remove aggregates, dimers, and monomers. The fractions were assessed on Blue Native-PAGE; pure-trimer fractions were pooled and stored at -80°C till further analysis. Protein concentration was determined by the bicinchoninic assay (BCA). PGT145-purification yields were 10-20% and PGT151-purification yields were 20-30% of the yield of Env obtained with 2G12 purification. Subsequent SEC yields were 30-40% trimer from 2G12-, 50-100% from PGT145-, and 70-100% from PGT151-purified Env.

### Field-flow fractionation (FFF) and Multi-Angle Light Scattering (MALS)

FFF experiments were performed with an Eclipse DualTec module (Wyatt Technology Corporation) running on an Agilent 1260 Infinity II platform (Agilent Technologies). Each sample (50 µg) was loaded onto a short channel unit assembled with a wide 350-µm spacer and a regenerated cellulose membrane (molecular-mass cutoff = 10 kDa). The detector flow was set to 0.4 ml/min, while cross-flow was 0.8 ml/min in all measurements. Total run length was 100 min.

The FFF system is coupled with an in-line MiniDawn Treos multiangle light scattering (MALS) detector, a quasi-elastic light scattering (QELS) detector, and an Optilab T-reX refractive index (RI) detector (Wyatt Technology Corporation). Molar mass and hydration radius (*R_h_*) values for different samples were determined based on the analysis of the light scattering and absorbance at UV 280 nm in Astra V software. These values were calculated only for the trimer-corresponding peak. Appropriate UV-280-nm extinction coefficients were applied for each sample. Refractive index increment (dn/dc) was set to 0.168, a value experimentally determined for fully glycosylated Env (137).

### Negative-stain electron microscopy (NS-EM)

Samples were diluted to ∼0.02 mg/mL final trimer concentration, and a 3-μl drop was applied to carbon-coated and plasma-cleaned copper mesh EM grids. The grids were stained with 2% (w/v) uranyl formate. Data were collected with an FEI Tecnai Spirit electron microscopy operating at 120 keV and equipped with a TVIPS TemCam F416 camera. Data processing steps, including particle picking, particle extraction, and 2D classification were performed with the Appion software suite (138). 2D class averages were analyzed and particles belonging to classes that did not unambiguously resemble native-like trimers (compact, trimeric structures) were categorized as ”non-native.”

### Cryo-electron microscopy (cryo-EM)

A mass of 0.2 mg of CZA97.012 SOSIP trimer (expressed in HEK-293F cells and purified by PGT151-affinity chromatography) was incubated with 0.3 mg of 3BNC117 Fab overnight at room temperature (approximately 2-fold molar excess of Fab per binding site calculated from the respective peptidic molecular masses of 215 and 50 kDa). On the following day the complex was purified on a gel-filtration column (HiLoad 16/600 Superdex 200 pg (Cytiva)), and the corresponding trimer:Fab complex peak was concentrated to ∼1 mg/ml. Immediately before freezing, Amphipol A8-35 (Anatrace) was added to a final concentration of 0.01% w/v to assist with particle tumbling. A 3-µl drop of the mixture was applied to a plasma-cleaned C-flat 2/2 holey carbon grid (Electron Microscopy Sciences), manually blotted and vitrified in liquid ethane with a homemade plunger.

Data were collected on a Thermo Fisher Titan Krios (300 keV) and a Gatan K2 Summit (4K × 4K) camera with Leginon software (139). The nominal magnification was 22500-fold resulting in a pixel size of 1.31 Å. A total of 1142 micrographs were collected in a defocus range of -1.0 to -2.7 µm and an average dose of 58 e-/Å^2^ per micrograph. Movies were aligned and dose-weighted with MotionCor2 (140) and imported into cryoSPARC (141). CTF corrections were performed with Gctf (142). Particles were picked with a combination of Blob Picker and Template Picker and subjected to rounds of 2D classification, Ab Initio 3D modeling, Heterogenous Refinement, and Non-Uniform Refinement (with global CTF corrections). C3 symmetry was applied to the final refinement of 79 261 particle images, resulting in an estimated ∼3.4-Å global FSC(0.143) resolution (**SI Table 3, SI Figure 5**).

Model building was initiated by generating a homology model of CZA97.012 with BG505 (PDB 4tvp) as the template and the UCSF Chimera (143) implementation of Modeller. Published 3BNC117 Fab models (PDB 4jpv) were fitted into the cryo-EM map. The complete model was then refined iteratively with Coot (144), Phenix real space refine (145), and Rosetta Relax (146). Final validation was performed with the MolProbity (147) and EMRinger (148) implementations in Phenix, and statistics are summarized in **SI Table 3**. The map and model have been deposited to the Electron Microscopy Data Bank and Protein Data Bank with accession codes summarized in **SI Table 3**.

### bNAb binding analyzed by ELISA

The binding of bNAbs to distinct clusters of epitopes on the CZA97.012 SOSIP.664 trimer expressed in HEK-293F was analyzed after 2G12, PGT145, or PGT151 purification followed by SEC was analyzed by ELISA as previously described (69, 83, 149). Briefly, 100 μl His-tagged trimer diluted to 1.5 μg/ml in PBS (pH 7.4) or PBS only as a control was added per well to Ni-NTA 96-well plates (Qiagen) and incubated over night at 4°C. After three washes with TBS, wells were blocked for 30 min at room temperature with 5% Milk (Bio-Rad 1706404) in PBS (100 μl/well) and again washed three times in PBS. The bNAbs were diluted in PBS with 5% Milk and 100 μl added per well. The plates were incubated for 2h at room temperature and then washed 3 times in TBS. Secondary HRP-conjugated antibody diluted 1/3000 in PBS with 10% Milk, 100 μl per well, was added. The plates were incubated for 45 min at room temperature and washed 5 times with PBS plus 0.05 % Tween. Then, 50 μl of TMB-substrate solution was added and the plates were incubated at room temperature for 3 minutes. To terminate the reaction, 50 μl of stop solution (0.3 N H_2_SO_4_) was added and the absorbance read at 450 nm on an Enspire multimode plate reader (Perkin Elmer). Background signals from control wells incubated with bNAbs and then conjugate were subtracted. Sigmoid functions were fitted to the data and curves plotted in Prism (GraphPad).

### Preparation of Fabs

3BNC117 and PGT151 Fabs were expressed by transient transfection of HEK-293F suspension cells with the respective *Fab* plasmids (36, 99). Fabs were affinity-purified on an anti-human-kappa-XL column, then subjected to ion-exchange fast protein liquid chromatography (ÄKTA FPLC, GE), and finally SEC. Fab purity was confirmed by reducing and non-reducing SDS-PAGE electrophoresis.

### bNAb binding analyzed by SPR

Antigenicity of bNAb epitopes on the His-tagged CZA97.012 SOSIP.664 trimer expressed in HEK- 293F cells and 2G12-, PGT145-, or PGT151- and then SEC-purified was also analyzed by SPR. Analyses were performed on BIAcore 3000 and T200 (Cytiva) instruments at 25^°^C as previously described (47, 99). Briefly, anti-His Ab covalently coupled to a CM5 sensor chip was used for capturing the His-tagged trimer to a density (*R_L_* value) of ∼250 RU. In each cycle, fresh trimer protein was captured, and at the end of each cycle the anti-His chip was regenerated with a single pulse of 10mM glycine (pH 2.0) for 1 min. The bNAb IgG was injected at a concentration of 500 nM; association was monitored for 300 s and dissociation for 600 s. Mass transfer limitation was avoided by a high flow rate (50 μl/min), which was used throughout also for regeneration to achieve better flushing. Sensorgrams were prepared in the BIAevaluation software.

Single-cycle kinetics (SCK on the T200 instrument only) was applied to the deep analysis of the binding of Fabs of PGT151 and 3BNC117 to differentially purified His-tagged CZA97.012 SOSIP.664 trimer. In each SCK experiment, increasing concentrations of Fabs (2-fold dilution starting from 1 μM) were injected sequentially in a single cycle. Fab at each concentration was allowed to bind for 300 s to anti-His-captured trimer on the sensor surface of CM5 chips. After the final injection, the dissociation was monitored for 3600 s. At the end of the cycle, the sensor surface was regenerated as above. In each SCK assay, 2 start-up cycles and 3 zero-analyte cycles with running buffer were included.

To consolidate the SCK results, the same Fabs were also analyzed by multi-cycle kinetics (MCK also on the T200 instrument). Fab solutions were titrated in 2-fold steps from 1 μM until no signal could be detected. Fab at each concentration was injected over freshly captured trimer; association was monitored for 300 s, and dissociation for 600 s. At the end of each dissociation phase, the sensor surfaces were regenerated, as above.

Trimer immobilization for the kinetic analyses was adjusted to give *R_max_* < 30 RU; experimental *R_max_* was ∼25 RU). A high flow rate (50 µl min^-1^) was used as a further measure to prevent mass transfer limitation. SCK and MCK data were analyzed with the BIAevaluation software v3.2.1 (Cytiva). Control-channel- and zero-analyte-subtracted sensorgrams were prepared. A Langmuir model was first fitted to the kinetic data. The Langmuir fit was considered acceptable based on goodness of fit (χ^2^<1), significance of kinetic rate constants (T>10), uniqueness (U<15, SCK only), and the scalar values of residuals (<2 RU). For PGT151 Fab, the Langmuir fit was unsatisfactory, and conformational-change and heterogeneous-ligand models were also fitted. They fulfilled the same criteria as the Langmuir model except that they do not include uniqueness evaluation. The conformational-change modeling was further investigated by injection-time variation: no deceleration in dissociation during 600 s was observed by reducing the association phase from 300 s to 30 s. The conformational-change modeling could therefore not be justified. Hence, the heterogeneous-ligand model was chosen and further validated (see Results). In all modeling, mass-transport limitation, counteracted by maximum flow rate and low trimer immobilization, was excluded by fulfillment of the following criteria: the mass-transfer constant, *k_t_,* was >10^8^ (RU. M^-1^. s^-1^) and its T value <10. As seen in SI Tables 1 and 2, these criteria were met with margins of several orders of magnitude.

### Global glycan analysis by HILIC-UPLC

The abundance of total N-linked glycans was determined after PNGase F treatment. PNGase F-released glycans were analyzed on a Waters Acquity H-Class UPLC instrument with a Glycan BEH Amide column (2.1 mm x 100 mm, 1.7 μm; Waters) and the following gradient: time (t) = 0: 22% A, 78% B (flow rate = 0.5 ml/min); t = 38.5: 44.1% A, 55.9% B (0.5 ml/min); t = 39.5: 100% A, 0% B (0.25 ml/min); t = 44.5: 100% A, 0% B (0.25 ml/min); t = 46.5: 22% A, 78% B (0.5 ml/min), where solvent A was 50 mM ammonium formate, pH 4.4 (Ludger) and B was acetonitrile. Fluorescence was measured at an excitation wavelength of 310 nm and an emission wavelength of 370 nm for procainamide, or 250 nm and 428 nm for 2-aminobenzoic acid (2-AA). Data were processed with Empower 3 software (Waters).

The relative abundance of each oligomannose-type glycan released by endoglycosidase H (Endo H), *i.e.*, Man_9_GlcNAc_2_, Man_8_GlcNAc_2_, Man_7_GlcNAc_2_, Man_6_GlcNAc_2_, and Man_5_GlcNAc_2_, was calculated by comparing with the total glycan pool released by PNGase F. An aliquot of 15 μl containing trimer with fluorescently labelled glycans was treated with Endo H overnight at 37°C (New England Biolabs). Glycans were extracted with a polyvinylidene-fluoride protein-binding membrane (Merck Millipore). Digested glycans were analyzed by HILIC-UPLC as before (51, 52). Glycans were quantified by integrating the peaks before and after glycosidase digestion, after normalization of the chromatograms.

### Site-specific glycopeptide analysis by LC-MS

The analyses were performed similarly to what has been described (92). Specifically, ∼50 μg of CZA97.012 SOSIP.664 trimer was denatured, reduced, and alkylated by sequential 1-hour incubations at room temperature as follows: 50 mM Tris-HCl buffer, pH 8.0, containing 6 M urea and 5 mM DTT, then in the dark after the addition of 20 mM iodoacetamide (IAA), and finally with an increased DTT concentration (20 mM) to eliminate residual IAA. The alkylated trimer was buffer-exchanged into 50 mM Tris-HCl, pH 8.0, in Vivaspin columns and aliquoted. Protein was digested with either trypsin or chymotrypsin (Mass Spectrometry Grade, Promega) at a mass ratio of 1:30 overnight at 37°C. Reaction mixtures were dried and glycopeptides were extracted with C18 ZipTip (Merck Millipore), following the manufacturer’s protocol.

Eluted glycopeptides were dried and re-suspended in 0.1% formic acid prior to mass spectrometric analysis. Aliquots of intact glycopeptides were analyzed by nano liquid-chromatography-electrospray-ionization mass spectrometry (LC-ESI MS) with an Easy-nLC 1200 system coupled to an Orbitrap Fusion mass spectrometer (Thermo Fisher Scientific) and fragmentation by higher-energy collisional dissociation (HCD). Peptides were separated on an EasySpray PepMap RSLC C18 column (75 μm x 75 cm) with a linear gradient consisting of 0% – 32% acetonitrile in 0.1% formic acid over 240 min followed by 35 min of 80% acetonitrile in 0.1% formic acid. The flow rate was set to 300 nl/min, the spray voltage to 2.5 kV, the temperature of the heated capillary to 275 °C, and HCD energy to 50%, appropriate for the fragmentation of glycopeptide ions. Data analysis and glycopeptide identification were performed with Byonic^™^ (Version 2.7) and Byologic^®^ software (Version 2.3; Protein Metrics Inc.), as follows. First, the mass spectrometry data and protein sequence were processed with Byonic^™^ allowing for a precursor mass tolerance of 4 ppm and a fragment mass tolerance of 10 ppm. Cleavage sites were specified as the C-terminus of Arg and Lys residues for tryptic digests, and the C-terminus of Tyr, Trp, and Phe residues for chymotryptic digests. The glycopeptide fragmentation data were then evaluated manually for each identified glycopeptide with Byologic^®^. The peptide was validated as true-positive when the correct b and y fragment ions were observed along with oxonium ions corresponding to the identified glycan structure. The intensities of the extracted ion chromatograms (XIC) over all charge states for each true-positive glycopeptide (with the same amino-acid sequence) were calculated as percentages, to determine the relative amounts of each glycoform.

To determine the class of glycan occupying particular sites, glycopeptides were digested with Endo H, which cleaves oligomannose- and hybrid-type glycans, leaving a single GlcNAc residue at the corresponding site. The reaction mixture was then dried completely and resuspended in a mixture containing 50 mM ammonium bicarbonate and PNGase F in pure O^18^- labelled water (Sigma-Aldrich) throughout. This second reaction cleaves the remaining complex-type glycans, leaving the GlcNAc residues intact. The use of H_2_O^18^ in this reaction enables complex glycan sites to be differentiated from unoccupied glycan sites as the hydrolysis of the glycosidic bond by PNGase F leaves an O^18^ isotope on the aspartate residue that results from the deamidation of the asparagine. The ensuing peptides were purified by C18 ZipTip, as outlined above and subjected to LC-ESI MS as described above but with a lower HCD energy of 27%, because glycan fragmentation was not required.

## ACKOWLEDGEMENTS

The human bNAbs VRC01, 2G12, PGT121, and PGT145 (68, 150–152) were graciously provided by the International AIDS Vaccine Initiative (IAVI, La Jolla), VRC34.01 (81) by John Mascola (VRC, NIH), 35O22 (153) by Mark Connors (VRC, NIH), and ASC202 by Marit J van Gils (AMC). Human Immunodeficiency Virus Type 1 (HIV-1) SG3ΔEnv Non-infectious Molecular Clone, ARP- 11051, contributed by Drs. John C. Kappes and Xiaoyun Wu (University of Alabama Birmingham (136)), was obtained through the NIH HIV Reagent Program, Division of AIDS, NIAID. We are grateful to Erik Francomano and Oscar Feliciano for expert technical assistance.

## DATA AVAILABILITY

The CZA97.012+3BNC117 cryo-EM map and model have been deposited to the Electron Microscopy Data Bank under accession code EMD-40088 and the Protein Data Bank under accession code 8gje, respectively.

